# Single-cell RNA sequencing uncovers cell-type-specific reprogramming of water-channel and cell-wall genes in PFOA uptake by lettuce root tips

**DOI:** 10.64898/2026.03.18.712360

**Authors:** Zhuo-Xing Shi, Peng-Fei Yu, Bo-Gui Pan, Yu-Qing Pu, Hai-Ming Zhao, Yan-Wen Li, Quan-Ying Cai, Nai-Xian Feng, Qing X. Li, Lei Xiang, Ce-Hui Mo

**Affiliations:** Guangdong Provincial Research Center for Environment Pollution Control and Remediation Materials, College of Life Science and Technology, Jinan University, Guangzhou 510632, China; State Key Laboratory of Ophthalmology, Zhongshan Ophthalmic Center, Sun Yat-sen University, Guangdong Provincial Key Laboratory of Ophthalmology and Visual Science, Guangdong Provincial Clinical Research Center for Ocular Diseases, Guangzhou 510060, China; Department of Molecular Biosciences and Bioengineering, University of Hawaii at Manoa, Honolulu, HI 96822, USA

**Keywords:** Crop absorption and accumulation, per- and polyfluoroalkyl substances (PFAS), root tip single-cell sequencing, cellular and molecular mechanism, food safety

## Abstract

Per- and polyfluoroalkyl substances (PFAS) are persistent organic pollutants that accumulate in edible crops, posing threats to food safety and human health. Breeding varieties with reduced PFAS accumulation offers a promising mitigation strategy, yet the underlying cellular mechanisms remain elusive. Here, we integrated short- and long-read single-cell RNA sequencing to generate a high-resolution transcriptomic atlas of lettuce (*Lactuca sativa L.*) root tips, comparing varieties with contrasting perfluorooctanoic acid (PFOA) accumulation capacities. Our analyses uncovered a synergistic mechanism underlying PFOA uptake: up-regulation of aquaporin genes in epidermal and xylem cells, coupled with down-regulation of cell-wall biosynthesis genes specifically in epidermal and xylem. Furthermore, we identified distinct RNA isoforms-including variants with altered coding potential-that may contribute to PFOA transport and deposition through structural modulation of the encoded proteins. This study represents the first application of single-cell sequencing to elucidate the cellular and molecular basis of crop responses to PFOA, providing new insights into pollutant-crop interactions and informing strategies for sustainable agriculture and food safety.

## Introduction

Perfluoroalkyl substances (PFASs) are an emerging class of persistent organic pollutants characterised by strong surface activity and exceptional chemical stability^1, 2^. These properties have driven their widespread use across industrial, agricultural, commercial, and domestic applications, leading to substantial and ongoing environmental emissions^3, 4^. Perfluorooctanoic acid (PFOA), a representative anionic PFAS with high water solubility, is frequently detected in agricultural soils, particularly in regions surrounding fluorochemical industrial facilities^5^. Its efficient uptake by crops and accumulation in edible tissues raise global concerns over food safety and present serious threats to ecosystem integrity and human health^5, 6, 7^. Toxicological and epidemiological evidences show that link chronic low-dose exposure to PFOA and related PFAS to nephrotoxicity, hepatotoxicity, reproductive toxicity and potential carcinogenicity^7, 8^. In recognition of its persistence and hazards, PFOA, its salts, and related compounds were listed as persistent organic pollutants (POPs) under Annex A of the Stockholm Convention in 2019. In 2020, the European Food Safety Authority (EFSA) set a total tolerable daily intake (TDI) of 0.63 ng kg⁻¹ day⁻¹ for combined exposure to PFOA, perfluorooctane sulfonate (PFOS), perfluorohexane sulfonic acid (PFHxS), and perfluorononanoic acid (PFNA), almost 2,400-fold lower than the TDI for PFOA alone established in 2008^6^. Many countries have since restricted or phased out PFOA^9^. Yet increasing evidences indicate that crop-derived dietary intake is a major human exposure route of PFOA because of its historical accumulation and persistence in soils^10, 11^, thereby underscoring the urgency of reducing its accumulation in edible plant parts.

Developing crop varieties with intrinsically low pollutant accumulation is an environmentally sustainable and cost-effective approach to safeguarding food quality^5^. Lettuce (*Lactuca sativa L.*), a globally cultivated leafy vegetable, serves as a valuable model. We identified low-accumulating varieties (LAVs) for PFOA, such as Italian curly lettuce, that retain 5-8 times less PFOA in edible tissues than high-accumulating varieties (HAVs), such as Qiangkun upright lettuce, under equivalent exposure conditions^5, 10^. Previous physiological studies have implicated differences in root uptake and translocation associated with exudation and transpiration rates contributed to these varietal disparities^10, 12, 13^. However, the cellular and molecular basis remains unclear, including the key cell types responsible for PFOA entry and transport and the regulatory genes that control these processes.

Single-cell RNA sequencing has transformed the ability to resolve transcriptional heterogeneity within complex plant tissues, enabling the identification of rare cell-types, regulatory circuits, and dynamic cell-cell interactions^14, 15^. In plants, such approaches have revealed the cellular complexity of root tips and their roles in environmental stress responses^16, 17, 18, 19^. Yet, the cell-type-specific processes and regulatory programs underlying organic pollutant uptake remain largely unexplored. For lettuce varieties with contrasting PFOA accumulation phenotypes, key questions remain: which root cell populations serve as the primary conduits for PFOA entry and transport? Which regulatory genes govern these processes, and how do their expression patterns differ across varieties and stress conditions? Addressing these questions requires a systematic, high-resolution dissection of root cell populations, integrating short- and long-read single-cell sequencing to resolve both gene expression and isoform diversity.

Here, we used an optimized protocol for protoplast isolation from lettuce (*Lactuca sativa L.*) root tips and applied both short-read and long-read single-cell RNA sequencing to profile root tip tissues. Leveraging an updated lettuce genome annotation from our previous work^20^, we substantially improved read mapping efficiency and data utility. We established a robust clustering and cell-type annotation framework, generating the first high-resolution single-cell atlas of lettuce root tips. A total of 66,156 high-quality single cells were profiled across four experimental conditions: high- (HAV) and low-accumulation (LAV) lettuce varieties, each with or without PFOA exposure. Our analyses revealed that up-regulation of aquaporin genes in epidermal and xylem cells, together with down-regulation of cell-wall biosynthesis genes specifically in the xylem, synergistically drives PFOA accumulation in lettuce roots. Using high-throughput long-read single-cell RNA sequencing (HIT-scISOseq)^21^, we further identified isoform-specific variants of aquaporin genes that are closely associated with PFOA uptake and deposition. Structural analyses indicate that these isoforms may exert regulatory effects without altering the encoded protein sequences. Together, these data provide a cell-resolved view of the mechanisms underlying varietal differences in PFOA accumulation and establish a foundation for breeding and management strategies aimed at reducing PFAS entry into the food chain.

## Result

### Study design and cell clustering

Previous surveys have documented widespread PFOA contamination in agricultural soils across China, with concentrations reaching 0.62 mg kg⁻¹. To reflect realistic pollution scenarios, we selected two lettuce (*Lactuca sativa L.*) varieties with contrasting PFOA accumulation capacities: HAV and LAV (Figure 1A). Plants were grown hydroponically for two weeks under PFOA stress at 0.1 mg L⁻¹ or 0.5 mg L⁻¹ (see Methods). After 12h of exposure, shoot tissues were harvested, and PFOA concentrations were quantified using a validated HPLC-MS/MS method. At 0.1 mg L⁻¹ PFOA, shoot PFOA concentrations in HAV were 3.33-fold higher than in LAV; at 0.5 mg L⁻¹, they were 2.16-fold higher (Figure 1B, Supplementary Table 1), confirming significant varietal differences in PFOA accumulation.

**Figure 1.**
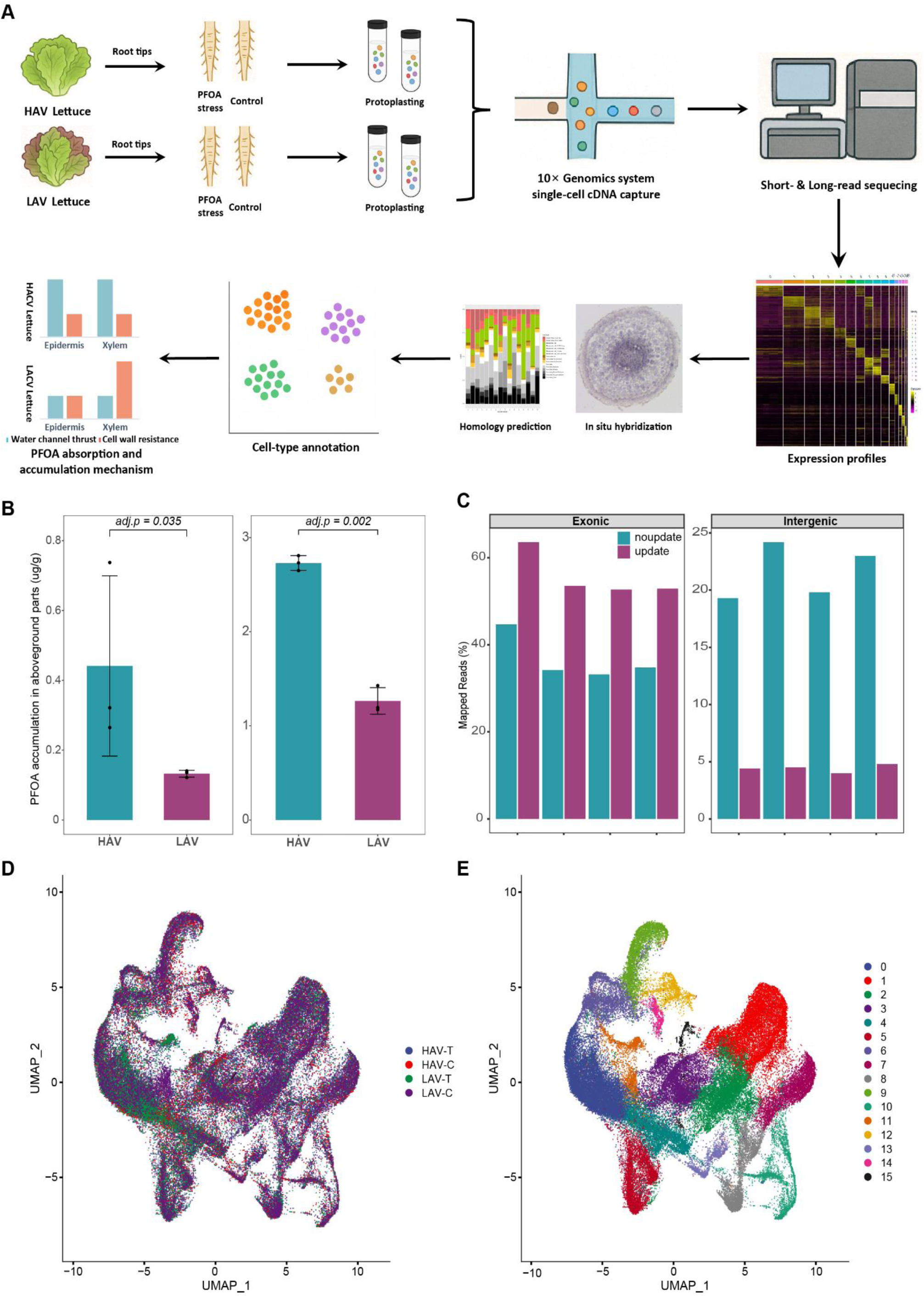
Single-cell sequencing of lettuce root tips reveals the cellular basis of differential PFOA accumulation. **A.** Schematic diagram of the experimental workflow: High-accumulating variety (HAV) and low-accumulating variety (LAV) of lettuces were subjected to PFOA stress. Root tips were collected and protoplasts isolated, followed by combined short-read and long-read single-cell sequencing. Cell clustering and annotation were performed to identify key PFOA-responsive genes. **B.** PFOA accumulation in edible parts of HAV and LAV lettuces under 0.1 mg·L⁻¹ and 0.5 mg·L⁻¹ PFOA stress. **C.** Proportion of single-cell sequencing reads mapped to exon and intergenic regions before and after gene annotation update. **D.** Uniform manifold approximation and projection (UMAP) visualization showing integration of single-cell data from four lettuce root tip samples. **E.** UMAP plot of cell clusters after integration, showing distribution of distinct cell populations.

We next performed 10×Genomics single-cell RNA sequencing, using both short-read and long-read approaches, on root tips of lettuce varieties under 0.5 mg L⁻¹ PFOA stress. Four parallel samples were collected: HAV without (HAV-C) and with (HAV-T) PFOA exposure, and LAV without (LAV-C) and with (LAV-T) exposure.

For short-read single-cell RNA sequencing, reads from the four samples were processed using CellRanger with both the pre-update and updated lettuce reference annotations. As shown in Supplementary Table 2, Figure 1C, and Supplementary Figure 1 the updated annotation markedly improved alignment performance: the confident genome alignment rate increased by 4.9-8.6%, confident exonic alignment increased by 18.1-18.9%, and confident intergenic alignment decreased by 14.9-19.7%. These improvements indicate that the updated annotation substantially increased the proportion of valid genome and exon alignments while reducing spurious intergenic alignments, thereby enhancing data utility and providing a more accurate foundation for downstream analyses.

Quality control of the short-read data yielded 80,849 high-quality single-cell profiles: 24,323 (HAV-T), 16,566 (HAV-C), 18,372 (LAV-T), and 21,588 (LAV-C). The average number of RNA molecules (UMIs) per cell ranged from 3,145 to 4,098, and the average number of detected genes ranged from 1,306 to 1,712, with a total of 26,977-27,485 unique genes detected across all cells (Supplementary Table 2-4).

Long-read single-cell RNA sequencing (HIT-scISOseq) produced raw CCS (circular consensus sequencing) data volumes of 22.28-29.14 GB per sample, with an average of 15-23 passes per CCS and mean quality values (QV) of 0.95-0.97. The number of full-length non-chimeric (FLNC) reads ranged from 20.12M to 23.96M (Supplementary Table 5-6), underscoring the high throughput and accuracy of the HIT-scISOseq platform.

Integrated analysis of all 80,849 cells using the Seurat IntegrateData function and clustree-guided resolution selection (Supplementary Figure 2A) yielded 16 transcriptionally distinct clusters (Cluster 0-15). Integration quality assessment demonstrated highly overlapping uniform manifold approximation and projection (UMAP) distributions across samples (Figure 1D-E, Supplementary Figure 2B, Supplementary Table 7), indicating that clustering reflected biological rather than technical variation. The final UMAP clustering revealed well-defined boundaries between the 16 clusters (Figure 1E), and sample-specific UMAPs confirmed consistent spatial organization across all four conditions (Figure 1D). These results validate the robustness of both the integration and clustering workflows, providing a solid framework for downstream cell-type annotation and functional analysis.

### Cell-type annotation of lettuce root tips

We first performed marker gene analysis for the 16 transcriptionally defined cell clusters using the FindAllMarkers function in Seurat, applying a significance threshold of p < 0.01 and an average log₂ fold-change (avg_log₂FC) > 0.6 (equivalent to ≥1.52-fold up-regulation). This analysis yielded 1,709 high-confidence marker genes. Heatmaps of the top 15 marker genes per cluster demonstrated highly specific expression patterns (Supplementary Figure 3A), while dot plots of the three most specific genes in each cluster highlighted both maximal expression levels and the proportion of expressing cells (Supplementary Figure 3B).

Given the absence of a comprehensive single-cell reference for lettuce roots, we leveraged eight *Arabidopsis thaliana* root single-cell datasets from scPlantDB (Supplementary Table 8) as annotation references. Cell-type classification was performed using SingleR (Figure 2A), in which each lettuce cell was compared against all eight *Arabidopsis* datasets and assigned the cell-type receiving the highest frequency of “votes”. The resulting annotation proportions are shown in Supplementary Figure 4 and Supplementary Figure 5A. This approach identified Cluster 12 and Cluster 14 as meristematic cells; Clusters 1-3 as ground tissue and root epidermis cells near the root cap-epidermis junction; Clusters 4-5 as pericycle cells; Clusters 7, 8, and 10 as phloem cells; Cluster 11 as root procambium cells; and Clusters 13 and 15 as xylem cells.

**Figure 2.**
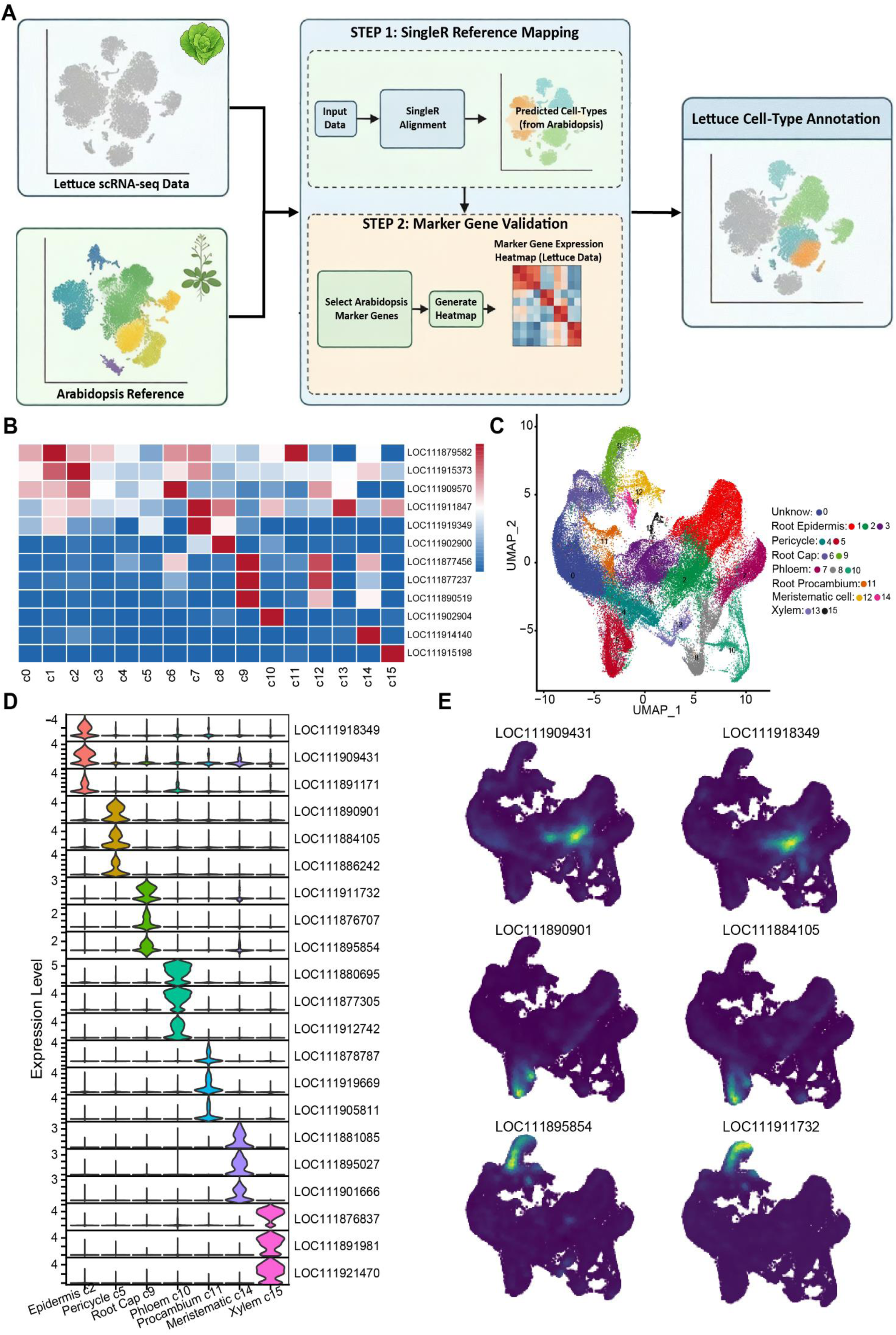
Cell type identification based on *Arabidopsis* reference data. **A.** Workflow of cell annotation: Two strategies, namely the SingleR method (leveraging an *Arabidopsis* single-cell dataset) and the use of classic *Arabidopsis* marker genes, were employed to assist in annotating and validating the lettuce single-cell data. **B.** Heatmap showing the expression patterns of classic *Arabidopsis* cell-type marker genes across distinct cell clusters in lettuce. **C.** Final cell type annotation of the single-cell atlas for lettuce root tips. **D-E.** Expression of cell-type-specific marker genes in lettuce root tips: (D) Violin plots. (E) Spatial expression feature plots.

To validate and refine these assignments, we extracted marker gene lists for each *Arabidopsis* root cell-type from scPlantDB and identified their lettuce homologs using OrthoFinder. We then quantified the concordance between lettuce cluster-specific markers and *Arabidopsis* homolog markers, considering both shared gene identities and overlapping GO terms. Bar charts in Supplementary Figure 5B-C illustrate a substantial fraction of markers and functional annotations overlapped, revealing high concordance between orthology-based analysis and SingleR-derived annotations.

We next examined expression patterns of conserved *Arabidopsis* cell-type markers in lettuce roots (Figure 2B, Supplementary Table 9). These included: SUC2 (stele-specific sucrose-proton symporter 2), ANAC033 (root cap-specific NAC domain protein 33), COBL9 (root hair-specific COBRA-like 9), ATPP2-A1 (phloem protein 2-A1), MYB36 (endodermis-specific Casparian strip regulator), RGF2 and PLT1 (meristematic regulators), IRX9 (xylem-specific irregular xylem 9), PDF1 and ATML1 (epidermis-specific transcription factors), LAX3 (lateral root-specific auxin transporter), and photosynthetic genes LHCA1 and LHCA2. These expression patterns were consistent with the SingleR-based annotations and provided additional resolution. In particular, *Arabidopsis* marker gene signatures supported Cluster 9 as root cap cells and suggested that Cluster 6 represents endodermal cells in a transitional state, spanning the root cap-epidermis boundary and stele differentiation zone.

One notable exception was Cluster 0, which exhibited markedly lower overall marker gene expression (Supplementary Figure 3) and yielded ambiguous SingleR assignments across Arabidopsis references (Supplementary Figure 4-5). We hypothesize that this cluster comprises low-quality cells. Following scPlantDB guidelines, we designated Cluster 0 as "Unknown" and excluded it from further analyses. After removal, the retained high-quality cell counts per sample were: HAV-T, 20,544; HAV-C, 13,374; LAV-T, 13,265; and LAV-C, 18,973, yielding a total of 66,156 cells for downstream investigations (Figure 2C, Supplementary Table 7). Violin plot and UMAP visualizations further illustrated the spatial distribution of markers across cell types, providing a functional and anatomical framework for downstream analyses (Figures 2D and 2E, Supplementary Figure 6).

Finally, we selected a subset of lettuce root tips marker genes (Supplementary Table 10) for spatial validation via RNA fluorescence in situ hybridization (RNA FISH). These included LOC111881417 (BT1 homolog), LOC111890901 (DETOXIFICATION 42 protein), LOC111914663 (uncharacterized proteins), LOC111880695 (PM19L membrane protein), LOC111878289 (histone H3.2), and LOC111876837 (cysteine protease XCP1). RNA FISH confirmed the strong, cell-type-specific expression patterns of these genes, fully consistent with our single-cell annotations and validating the robustness of the cell-type classification framework (Supplementary Figure 7).

### Trajectory analysis of cell-types in lettuce root tips

We performed Monocle2 pseudotime analysis on four cell groups with distinct differentiation characteristics: Cluster 8 (phloem), Cluster 11 (root procambium), and Clusters 12 and 14 (meristematic cells). The analysis resolved five discrete developmental states, which coalesced into two major developmental nodes (Figure 3A). State 1, dominated by meristematic cells from Clusters 12 and 14, represents a highly proliferative phase. State 2 captures the early transition of meristematic cells toward root procambium (Cluster 11) and phloem (Cluster 8), marking the onset of tissue-specific differentiation. State 4 reflects the direct differentiation of Cluster 12 into root procambium, signifying a pivotal shift from meristematic to root cell identity. State 3 delineates a more complex trajectory, in which Clusters 12 and 14 first differentiate into root procambium (Cluster 11) before further commitment to phloem (Cluster 8). State 5 comprises undifferentiated meristematic cells within Cluster 14. Together, these trajectories outline a developmental progression, namely, Cluster 14 → Cluster 12 → Cluster 11 → Cluster 8, that aligns with established root apex developmental hierarchies (Figure 3B).

**Figure 3.**
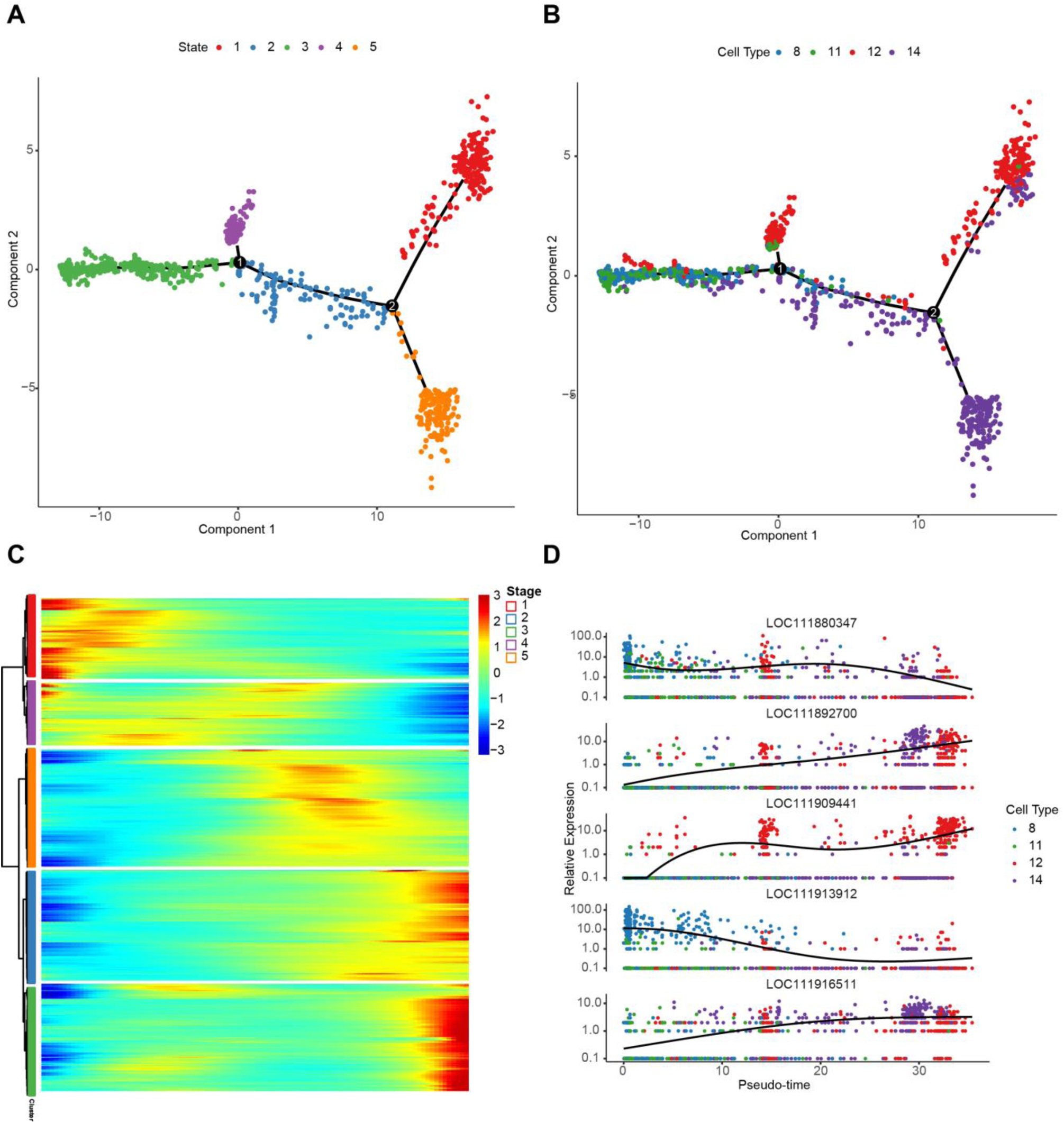
Single-cell developmental trajectory analysis of lettuce root tips. **A-B.** Trajectory analysis reveals the developmental paths of lettuce root tip cells from meristematic cells towards procambium and phloem differentiation. **C.** Heatmap showing the dynamic expression trends of stage-specific genes along the pseudotime developmental trajectory in lettuce root tips. **D.** Scatter plot illustrating the expression dynamics of stage-specific genes across various developmental stages along the pseudotime trajectory in lettuce root tips.

Interpretation of these trajectories suggests that Cluster 14 constitutes the stem cell niche of the meristematic zone, enriched in highly proliferative, putative stem cells, whereas Cluster 12 represents a differentiation-active domain, containing cells transitioning toward specialized fates.

The expression heatmap in Figure 3C depicts state-specific genes along pseudotime, with pseudotime on the x-axis, genes on the y-axis, and a color gradient reflecting relative expression. Genes enriched in State 1 peak early in pseudotime, while State 3 genes exhibit late-stage expression. These coordinated temporal patterns reveal modules of co-regulated genes acting at similar developmental windows, providing mechanistic insights into stage-specific transcriptional programs. Figure 3D and Supplementary Table 11 summarizes statistical metrics for state-specific genes and highlights representative loci that may regulate root apex cell fate transitions.

### Identification of key PFOA-responsive genes at single-cell resolution in lettuce root tips

We performed differential expression gene (DEG) analysis across cell clusters 1-15 using the FindMarkers function in Seurat, focusing on two pairwise comparisons: HAV-C vs. LAV-C and HAV-T vs. LAV-T, to assess varietal differences and stress-induced transcriptional changes. For each cluster, candidate DEGs were retained based on stringent criteria: (1) p < 0.05; (2) pct1 > 0.2 for up-regulated genes, to exclude sparsely expressed transcripts; (3) |avg_log_₂_FC| > 1, ensuring at least a twofold change in expression; and (4) |avg_log_₂_FC| difference > 0.5 relative to the HAV-C vs. LAV-C baseline, to control for constitutive varietal effects. This filtering yielded 3,397 DEGs across all clusters (Figure 4A, Supplementary Table 12).

**Figure 4.**
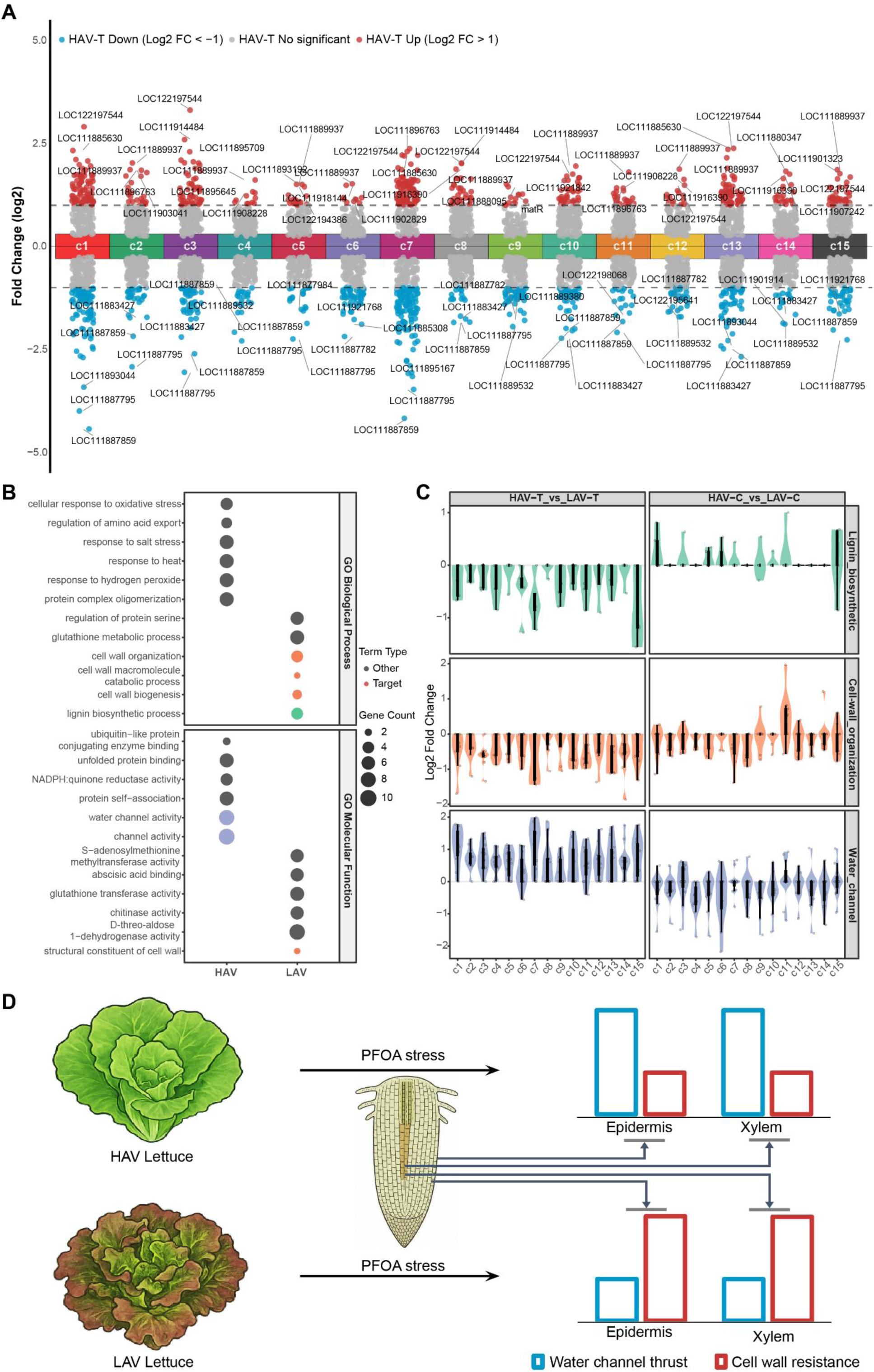
Analysis of cell type-specific differentially expressed genes in response to PFOA in root tips of high- and low-accumulation lettuce varieties. **A.** Scatter plot showing variety-specific differentially expressed genes across cell-types after PFOA treatment. Red and blue dots represent genes significantly up-regulated in HAV and LAV, respectively. **B.** GO enrichment analysis of selected variety-specific PFOA-responsive differentially expressed genes. Pathways of focus-aquaporin activity, cell-wall synthesis, and lignin synthesis-are highlighted in red. **C.** Single-cell expression distribution of key genes related to aquaporins, cell-wall synthesis, and lignin synthesis in epidermal and xylem cells of lettuce root tips. **D.** Schematic diagram illustrating the dynamic changes in aquaporin function and cell wall structure in root tips of HAV and LAV lettuce under PFOA stress.

After merging results and removing duplicates, we obtained a non-redundant set of 468 genes, comprising 206 genes up-regulated in HAV-T and 262 genes up-regulated in LAV-T. Functional annotation using an updated database revealed distinct transcriptional signatures (Figure 4B): genes up-regulated in HAV-T were significantly enriched for water-channel activity, whereas those up-regulated in LAV-T were enriched for cell-wall organization and lignin biosynthetic processes.

Within the aquaporin-enriched set, several genes displayed strong cluster-specific regulation under stress (Figure 4C, Supplementary Figure 8). LOC111909461, LOC111901322, and LOC111901323 were markedly up-regulated in clusters 1, 2, 3, and 15 of HAV-T. For cell-wall-biosynthesis-related genes, several exhibited coordinated down-regulation in HAV-T. LOC111909624 was suppressed in cluster 2 of both HAV-T and HAV-C. LOC111880393, LOC111887823, and LOC111880378 were down-regulated in clusters 1 and 7 of HAV-T; notably, LOC111887823 was also suppressed in both HAV-T and HAV-C in these clusters. LOC111901914 and LOC111899844 were specifically down-regulated in cluster 11 of HAV-T, and LOC111888139 in cluster 15 of HAV-T. These patterns suggest an inhibition of xylem cell wall formation in HAV-T. In the lignin biosynthesis pathway, LOC111876731 and LOC111899992 were down-regulated in cluster 15 of HAV-T.

Together, these results support a model in which varietal differences in PFOA accumulation are driven by coordinated regulation of aquaporin-mediated transport and xylem cell wall architecture (Figure 4D). Under PFOA stress, HAV exhibited increased aquaporin expression in epidermal and xylem tissues, consistent with enhanced PFOA uptake. Concurrently, reduced expression of cell-wall-biosynthetic genes likely weakens the adhesive retention of PFOA within the xylem, facilitating its transport. In contrast, LAV down-regulated aquaporins and up-regulated cell-wall-synthesis genes, which may limit PFOA entry and reinforce its retention in xylem tissues. This antagonistic transcriptional program provides a mechanistic basis for variety-specific responses to PFOA exposure by jointly modulating transport capacity and structural retention in root tissues.

### Identification of key PFOA-responsive RNA isoforms in lettuce root tips

To integrate the HIT-scISOseq dataset with an existing short-read single-cell reference atlas, we transferred both cell coordinates and cluster identities from the atlas to the HIT-scISOseq data. UMAP projections of 16 clusters across four samples (Figures 5A-B, Supplementary Figure 9) revealed highly concordant cluster distributions with sharply defined boundaries, underscoring the robustness of HIT-scISOseq for high-resolution single-cell analysis.

**Figure 5.**
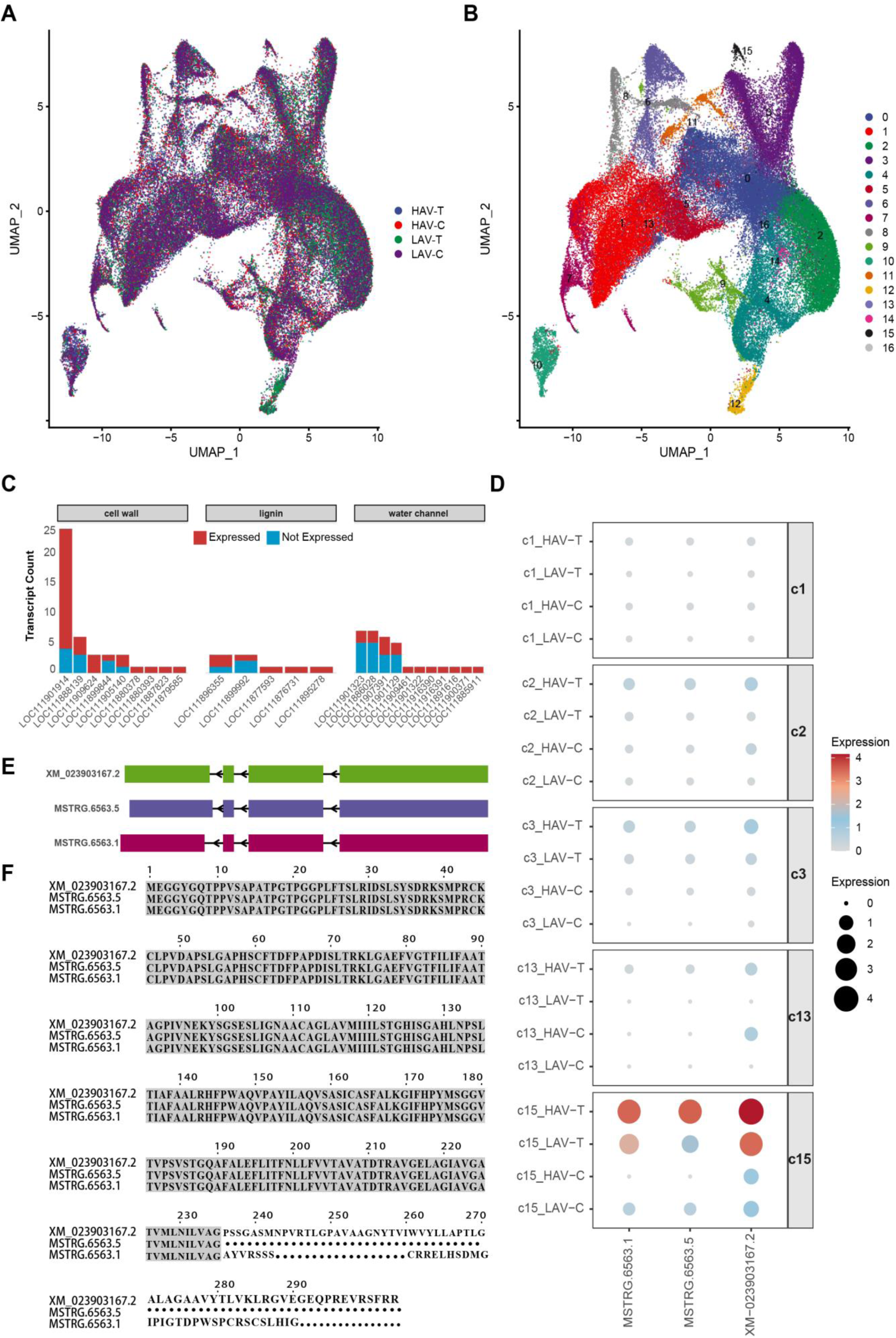
Analysis of differential RNA isoform usage patterns of aquaporin genes in lettuce root tip xylem using HIT-scISOseq. **A.** UMAP visualization showing the integration of four samples based on the RNA isoform expression matrix derived from HIT-scISOseq. **B.** UMAP plot of cell clusters based on the RNA isoform expression matrix. **C.** Bar plot showing expression levels of RNA isoforms for genes involved in cell wall synthesis, lignin synthesis, and aquaporin activity in the HIT-scISOseq dataset. **D.** Dot plot displaying expression differences of three detected RNA isoforms of the aquaporin gene LOC111907391 in epidermal versus xylem cells. **E.** Exon structure schematic of the three expressed RNA isoforms of LOC111907391. **F.** Protein sequence alignment of the three expressed RNA isoforms of LOC111907391.

Using Seurat’s FindAllMarkers function, we identified cluster-specific transcript isoforms in the HIT-scISOseq dataset. Heatmaps of the top 15 marker transcripts per cluster (Supplementary Figure 10) demonstrated strong cluster specificity, validating the accuracy and reproducibility of isoform-level clustering.

We next quantified the expression of transcripts associated with water-channel, cell-wall biosynthesis, and lignin biosynthesis. Among 11 genes annotated with multiple reference transcripts, nine expressed multiple isoforms in the HIT-scISOseq data (Figure 5C, Supplementary Table 13). In total, we identified 55 expressed isoforms, of which 25 were from the reference annotation, 22 were from our updated gene annotation, and 8 were novel isoforms discovered through HIT-scISOseq, highlighting the value of our dataset in uncovering transcriptomic complexity (Supplementary Figure 11). Notably, three isoforms of the aquaporin gene LOC111907391 (XM_023903167.2, MSTRG.6563.5, and MSTRG.6563.1) were significantly up-regulated in Cluster 15 of HACV-T samples (Figure 5D). This finding reveals a striking cell-type-specific isoform expression pattern.

Exon-structure comparison showed that these isoforms differ primarily at the 3′ end of exon 4, potentially altering protein translation (Figure 5E). Open reading frame (ORF) prediction indicated that MSTRG.6563.5 and MSTRG.6563.1 encodes a truncated protein (Figure 5F). Such divergence may affect pore selectivity, and functional responses to PFOA stress in lettuce root tip cells.

### Structural characterization and molecular docking of aquaporin isoforms

To probe structural and functional implications, we predicted the 3D conformations of the three isoforms using AlphaFold2. All exhibited canonical aquaporin architectures with α-helical transmembrane domains (Supplementary Figure 12), yet showed pronounced differences in the disordered C-terminal regions. These regions could modulate protein-protein interactions, subcellular localization, and stability.

To elucidate the molecular mechanisms underlying PFOA uptake and transport in lettuce (*Lactuca sativa L.*) under PFOA stress, we performed structural modeling analyses on three aquaporin isoforms (XM_023903167.2, MSTRG.6563.5, and MSTRG.6563.1) encoded by LOC111907391, a NIP5-1 aquaporin gene identified in root tips (Figure 6). Channel profiling and pore diameter calculations using the HOLE program revealed distinct spatial topologies between the full-length and truncated protein isoforms. The full-length protein XM_023903167.2 exhibited a relatively narrow channel opening (Figures 6A and 6D), whereas the two truncated isoforms, MSTRG.6563.5 and MSTRG.6563.1, displayed wider pore apertures (Figures 6B-C and 6E-F). This pore expansion is expected to enhance transport capacity, providing a structural basis for more efficient translocation in plants. Consistent with these structural findings, transcriptomic data revealed that LOC111907391 was significantly induced in the xylem of high-accumulation (HAV) lettuce under PFOA stress. Together, the expression and modeling data support a model in which PFOA exposure promotes the expression of truncated aquaporin isoforms with wider channels (MSTRG.6563.5 and MSTRG.6563.1). These isoforms may partially substitute for, or act alongside, the full-length protein to reduce transport resistance and enhance PFOA uptake and xylem translocation. This finding provides important molecular insights into varietal differences in organic pollutant accumulation in plants.

**Figure 6.**
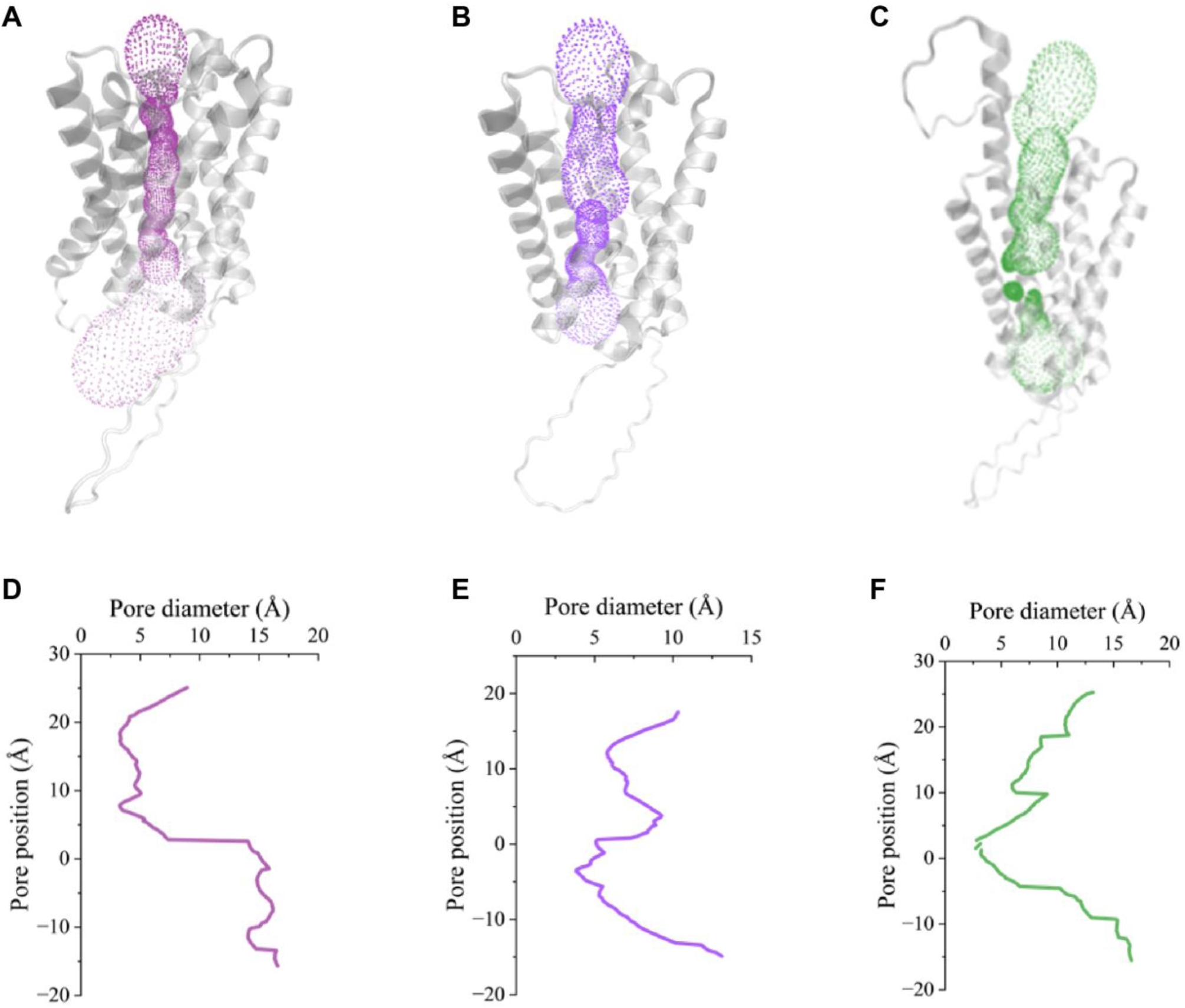
Structural characterization analysis of aquaporin LOC111907391 isoforms. **A-B.** Visualization of the internal channel profiles for XM_023903167 (full-length), MSTRG_6563_5 (truncated), and MSTRG_6563_1 (truncated), respectively. **D-F.** Quantitative distribution of channel pore diameters corresponding to the three isoforms, illustrating the widened apertures in truncated versions. Pore diameter calculations were performed using the HOLE program. Protein structures and stable binding configurations were visualized and rendered using PyMOL.

## Discussion

This study provides the first single-cell transcriptomic dissection of the cellular and molecular processes governing perfluorooctanoic acid (PFOA) uptake and accumulation in lettuce roots. By integrating short-read and long-read sequencing, we generated a high-resolution cellular atlas and resolved the gene- and isoform-level programs that distinguish high- from low-accumulating lettuce varieties. Our findings reveal a coordinated mechanism in which epidermal aquaporins and xylem cell-wall architecture jointly determine the efficiency of PFOA transport, offering new entry points for mitigating PFAS contamination in crops.

The root epidermis and xylem emerged as pivotal control points for PFOA uptake and translocation. In high-accumulating variety (HAV), epidermal cells exhibited pronounced up-regulation of aquaporin genes (e.g., LOC111907391, LOC111901322), consistent with the capacity of certain water-channel proteins to mediate passive transport of small polar contaminants, including PFAS. In parallel, reduced expression of cell wall biosynthetic genes (e.g., LOC111909624, LOC111880393) in xylem cells suggests diminished lignification and wall rigidity, conditions that may lower apoplastic resistance and enhance pollutant movement into vascular tissues^10^. Low-accumulating variety (LAV) displayed the inverse pattern, with suppressed aquaporin activity and reinforced cell wall synthesis, forming a dual barrier against PFOA entry and vascular loading. These cell-type-specific transcriptional signatures highlight the fine spatial regulation underlying pollutant accumulation phenotypes.

Long-read single-cell sequencing (HIT-scISOseq) uncovered an additional regulatory tier: PFOA-induced generation of functionally distinct RNA isoforms. For the aquaporin LOC111907391, alternative splicing produced C-terminal variants predicted to differ in stability and solute selectivity, echoing evidence from *Arabidopsis* where isoform diversity modulates aquaporin trafficking and stress adaptation. In contrast, LOC111909624 isoforms encoded identical proteins, yet their cell-type-specific abundance points to transcriptional fine-tuning of cell wall remodeling under PFOA exposure. Such isoform-level regulation is rarely considered in pollutant uptake studies but may be critical for environmental responsiveness.

Pseudotime analysis reconstructed trajectories from meristematic cells to procambium and phloem lineages, revealing stage-specific engagement of PFOA-responsive genes In LAV, differentiating xylem cells were enriched for lignin biosynthetic transcripts, suggesting that early reinforcement of secondary walls physically restricts pollutant mobility, naemly, a mechanism reminiscent of lignification-based metal sequestration in hyperaccumulator species. These results imply that modifying developmental programs could be an underexplored strategy for engineering low-accumulating crops.

The cell-type- and isoform-specific regulation of LOC111907391 and LOC111909624 makes them strong candidates for functional validation. Suppressing aquaporin activity in HAV or enhancing cell wall synthesis in LAV could shift accumulation profiles toward food safety thresholds. The conservation of orthologous regulators (e.g., SUC2, IRX9) across species suggests that these principles may generalize to other edible crops. In parallel, high-accumulating varieties could could support phytoremediation of PFAS-contaminated soils, while low-accumulating lines safeguard production in affected areas.

This study emphasizes transcriptional control and does not address post-translational regulation, such as phosphorylation-driven gating of aquaporins, or the potential influence of rhizosphere microbiomes on PFAS bioavailability. Integrating single-cell proteomics and metabolomics could address these layers. Field trials will also be essential to determine the stability of low-accumulation traits under agronomic conditions. Expanding this framework to other PFAS and diverse crop species will be critical for managing persistent organic pollutants in agriculture.

By resolving the root tip at single-cell resolution, we delineate how the interplay between water-channel and cell-wall architecture regulation PFOA accumulation. This integrated view offers both mechanistic insight and actionable targets for breeding programs, supporting the development of crops that are productive and intrinsically resilient to chemical contamination.

## Methods

### Hydroponic cultivation of lettuce

Seeds of two lettuce varieties (upright and purple-leaf) were surface-sterilized with 10% H₂O₂ for 30 min, rinsed thoroughly with deionized water, and air-dried. Seeds were then uniformly sown onto vermiculite seedling trays. When seedlings developed three true leaves, those with uniform growth were selected, their roots washed with deionized water, and transferred to a hydroponic system. Each hydroponic unit contained 2 L of nutrient solution, with pH maintained at 6.0-6.2. Aeration was provided via an air pump for the initial 24 h. Two PFOA (perfluorooctanoic acid) stress concentrations (0 mg/L and 0.5 mg/L) were applied to both lettuce varieties. After 12 h of PFOA treatment, aboveground tissues were harvested, and PFOA content was quantified using high-performance liquid chromatography-tandem mass spectrometry (HPLC-MS/MS).

### Root tip protoplast isolation

Healthy lettuce root tips were excised into 1-cm segments for protoplast isolation. The procedure was as follows:

1. Root segments were fully submerged in pre-prepared enzymatic digestion solution in a sterile 50-mL conical flask.
2. The flask was subjected to vacuum infiltration for 5–30 min in darkness until no bubbles were observed.
3. Enzymatic digestion was performed at 40 rpm on a shaker in darkness for 3-4 h.
4. An equal volume of W5 solution was added to stop the enzymatic reaction, followed by gentle mixing.
5. The mixture was filtered through a 200-mesh sieve pre-wetted with W5 solution into a 50-mL round-bottom centrifuge tube.
6. Centrifugation was performed at 100 g for 8 min at 4°C (acceleration/deceleration set to 3-4).
7. The supernatant was carefully removed, and protoplasts were gently resuspended in 5 mL pre-chilled W5 solution, followed by a second centrifugation at 100 g for 8 min at 4°C.
8. The supernatant was discarded, and protoplasts were resuspended in an appropriate volume (∼1 mL) of W5 solution, incubated on ice for 30 min, and stored for downstream use.

### Short-read single-cell RNA sequencing of lettuce root tips

The single-cell sequencing workflow was as follows:

1. Protoplasts were washed and resuspended in PBS containing 0.04% BSA, then filtered through a 30-40 μm cell strainer to remove large cell clumps or debris, preventing microfluidic chip clogging.
2. Cell viability was assessed using Trypan Blue exclusion, and cell counts were quantified with an automated cell counter to determine the ratio of live to dead cells.
3. Single-cell dissociation was performed using the 10× Genomics Chromium Single Cell 3’ GEM, Library, and Gel Bead Kit v3 (PN-1000075) on the GemCode platform, targeting recovery of ∼20,000 cells.
4. Illumina libraries were prepared following the Chromium Single Cell 3’ Reagent Kits User Guide, V3 Chemistry.
5. Sequencing was conducted on the Illumina NextSeq 550 system (SY-415-1002) using NextSeq High Output Kits (150 cycles; 20024907) with the following parameters: Read1, 150 cycles; I7 index, 8 cycles; Read2, 150 cycles.
6. Sequencing data were processed using CellRanger software with an updated GTF file to generate a single-cell gene expression matrix.

### Long-read single-cell RNA sequencing of lettuce root tips

The long-read single-cell sequencing workflow was as follows:

1. After completing step 2.1 of the 10× Genomics Single Cell 3’ RNA library preparation (full-length cDNA synthesis), half of the purified cDNA was used for PacBio library construction.
2. Full-length cDNA products were processed using the BGI RNA ligation kit for second-strand cDNA synthesis, artifact removal, RNA ligation, and HIT-scISOseq library construction.
3. Libraries were sequenced on the PacBio Sequel II system.
4. Sequencing data were analyzed using scISA-tools (https://github.com/shizhuoxing/scISA-Tools) for circular consensus sequence (CCS) deconvolution, full-length non-concatemer (FLNC) read extraction, and cell barcode correction. FLNC reads were aligned to the lettuce reference genome using minimap2, and isoform identification and quantification were performed with IsoQuant using an updated GTF file, yielding a single-cell isoform expression matrix.

### Cell clustering analysis

Single-cell data from four lettuce root tip samples (80,849 cells) were integrated using the IntegrateData function in Seurat^22^ to correct for batch effects. The impact of different resolution parameters (0-1, step size 0.1) on cell clustering was systematically evaluated using clustree^23^. Ultimately, the 80,849 cells were classified into 16 clusters (Cluster0-Cluster15).

### Cell-type annotation

Cell type annotation was performed using SingleR^24^ by comparing the gene expression profiles of lettuce root tip single cells with annotated Arabidopsis datasets in the scPlantDB^25^ database. The most frequently assigned cell type was selected as the final annotation for each cell to minimize dataset-specific biases.

Candidate marker genes for each cell cluster were identified using Seurat’s FindAllMarkers function. Orthologous genes between lettuce and Arabidopsis were identified using OrthoFinder^26^. Marker gene sets for lettuce clusters were compared with Arabidopsis root tip marker gene lists from scPlantDB, and consistency was evaluated by calculating the percentage of shared genes, visualized as bar plots. The results were highly consistent with SingleR annotations.

Conserved Arabidopsis cell-type-specific marker genes were identified in lettuce via ortholog mapping, and their expression patterns across lettuce root tip clusters were analyzed. These results corroborated SingleR annotations. Notably, Cluster9 was further subdivided into root cap cells, and Cluster6 was identified as potentially representing endodermal cells in the root cap-epidermis to stele differentiation zone, providing insights into lettuce root tip cell differentiation.

To validate SingleR annotations, RNA fluorescence in situ hybridization (RNA FISH) probes were designed for cluster-specific marker genes. RNA FISH experiments confirmed the spatial localization of these markers, enhancing the reliability of bioinformatic annotations.

### Single-cell developmental trajectory analysis

The Monocle2^27^ algorithm was used to infer cell differentiation trajectories. This unsupervised method employs reverse graph embedding to position cells along developmental trajectories based on single-cell transcriptomic data, enabling pseudotime analysis to elucidate cell differentiation or subtype evolution and identify key regulatory genes.

### Differential gene and isoform expression analysis

Differentially expressed genes (DEGs) and RNA isoforms across cell clusters were identified using Seurat’s FindMarkers function. Gene Ontology (GO) and KEGG pathway enrichment analyses were performed using clusterProfiler^28^. Statistical analyses were conducted in R, with data visualization generated using ggplot2.

### Aquaporin structure prediction and PFOA interaction analysis

Exon structures of RNA isoforms were visualized using the ggtranscript^29^ R package. Three aquaporin isoforms (XM-023903167.2, MSTRG.6563.5, and MSTRG.6563.1) were subjected to three-dimensional structure prediction using AlphaFold2^30^, channel pore analysis used HOLE^31^, and structural comparisons were performed and visualized with PyMOL.

## Data availability

The HIT-ISOseq sequencing data and updated annotation file are available at NCBI Gene Expression Omnibus (https://www.ncbi.nlm.nih.gov/geo/) under the accession numbers GSE264560. The reference genome and gene annotation file (Lsat_Salinas_v7) were downloaded from NCBI. All single-cell data will be officially made public before the papers are accepted. All other data are available from the corresponding author upon reasonable request.

## Acknowledgements

This work received support from the National Natural Science Foundation of China (no. 42030713 to C.-H.M.; no. 42477222, 42177187, 42077300 to L.X.; and no. 42107148 to Z.-X.S., 42377236 to Y.-W.L).

## Author Contributions Statement

C.-H.M, L.X., and Z.-X.S. conceived and designed the project; Z.-X.S., L.X., and P.-F.Y. developed the experimental technology; Y.-Q.P., B.-G.P., and P.-F.Y. performed sequencing experiments. Z.-X.S., Y.-Q.P., and P.-F.Y. performed the informatics analysis; Z.-X.S., Y.-Q.P., and P.-F.Y. coordinated data release and assisted with executing the pipeline. C.-H.M., L.X., Z.-X.S., Y.-Q.P., P.-F.Y., B.-G.P., H.-M.Z., N.-X.F., Q.-Y.C., Y.-W.L., and Q.-X.L. wrote the manuscript and created the figures; H.-M.Z., N.-X.F., Q.-Y.C., and Y.-W.L. advised the study and revised the manuscript. All authors have read and approved the final version of this manuscript.

