## Supplementary Figure1-12 and Supplementary Table1-13 for "Single-cell RNA sequencing uncovers cell-type-specific reprogramming of water-channel and cell-wall genes in PFOA uptake by lettuce root tips"

### Supplementary Information

#### TABLE OF CONTENTS

- **Supplementary Figures**

Supplementary Figure 1. CellRanger processing statistics of raw scRNA-seq data.
Supplementary Figure 2. Resolution optimization and clustering analysis of integrated single-cell data from four samples.
Supplementary Figure 3. Identification and expression of cluster marker genes.
Supplementary Figure 4. Bar plot showing the results of lettuce root tip single-cell annotation using eight Arabidopsis data.
Supplementary Figure 5. Bar plots shows the annotation results of lettuce root tip single-cell data based on three strategies.
Supplementary Figure 6. FeaturePlot visualization of cell type-specific marker genes.
Supplementary Figure 7. RNA fluorescence in situ hybridization validation of selected cell type-specific marker genes.
Supplementary Figure 8. DotPlot visualization of water-channel, cell-wall, and lignin biosynthesis gene expression.
Supplementary Figure 9. UMAP plot of HIT-scISOseq cell clusters after integration of four samples.
Supplementary Figure 10. Heatmap of top 15 marker isoform for each cell cluster based on the integrated single-cell data.
Supplementary Figure 11. Expression profiles of RNA isoforms from genes in the HIT-scISOseq dataset.
Supplementary Figure 12. Predicted 3D structures of three proteins encoded by the aquaporin gene LOC111907391.

- **Supplementary Tables**

Supplementary Table 1. Statistics of PFOA accumulation in two lettuce varieties.
Supplementary Table 2. Statistics of single-cell NGS sequencing data.
Supplementary Table 3. Statistics on the quality control of expression matrices from NGS sequencing.
Supplementary Table 4. Detection statistics of RNA molecules and isoforms in NGS matrix.
Supplementary Table 5. Statistics of HIT-scISOseq sequencing data from lettuce root tips.
Supplementary Table 6. Detection statistics of RNA molecules and isoforms in HIT-scISOseq matrix.
Supplementary Table 7. Statistics of cell numbers in each cell cluster.
Supplementary Table 8. scPlantDB datasets used for cell-type annotation in lettuce root tip.
Supplementary Table 9. Homologs of conserved Arabidopsis marker genes in lettuce and their associated cell types.
Supplementary Table 10. RNA FISH probe sequence.
Supplementary Table 11. Differentially expressed genes in distinct states along the developmental trajectory.
Supplementary Table 12. Differential gene expression results among lettuce root tip cell clusters.
Supplementary Table 13. Transcript expression statistics of key genes involved in PFOA accumulation.

#### ■ Supplementary Figures

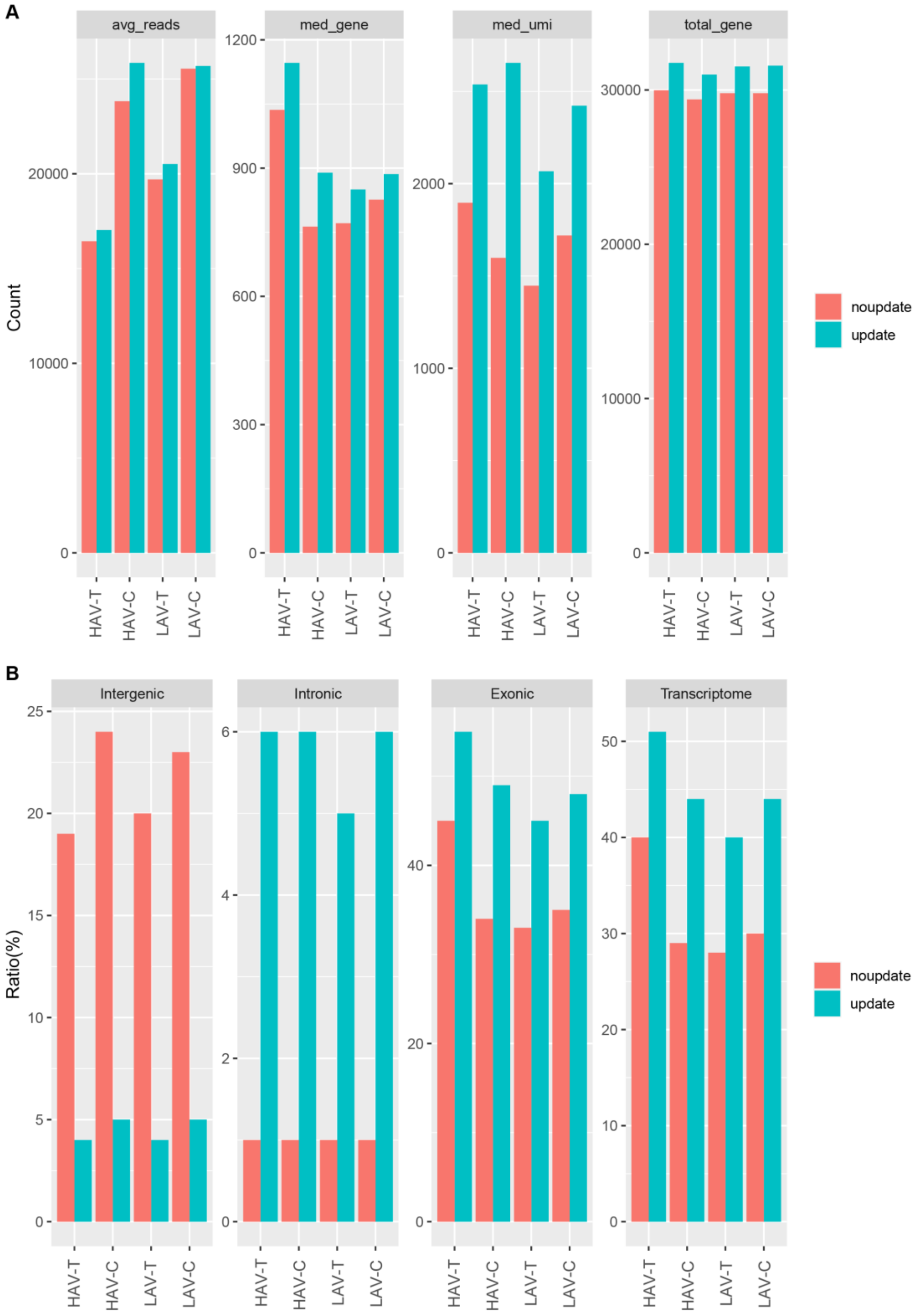


**Supplementary Figure 1. CellRanger processing statistics of raw scRNA-seq data. A.** Bar plots of per-cell metrics for each sample: average number of reads (avg_reads), median number of detected genes per cell (med_gene), median number of UMIs per cell (med_umi), and total number of detected genes (total_gene). **B.** Bar plots showing the percentage of reads mapped to different genomic regions per sample at single-cell level: intergenic, intronic, exonic, and transcriptome.

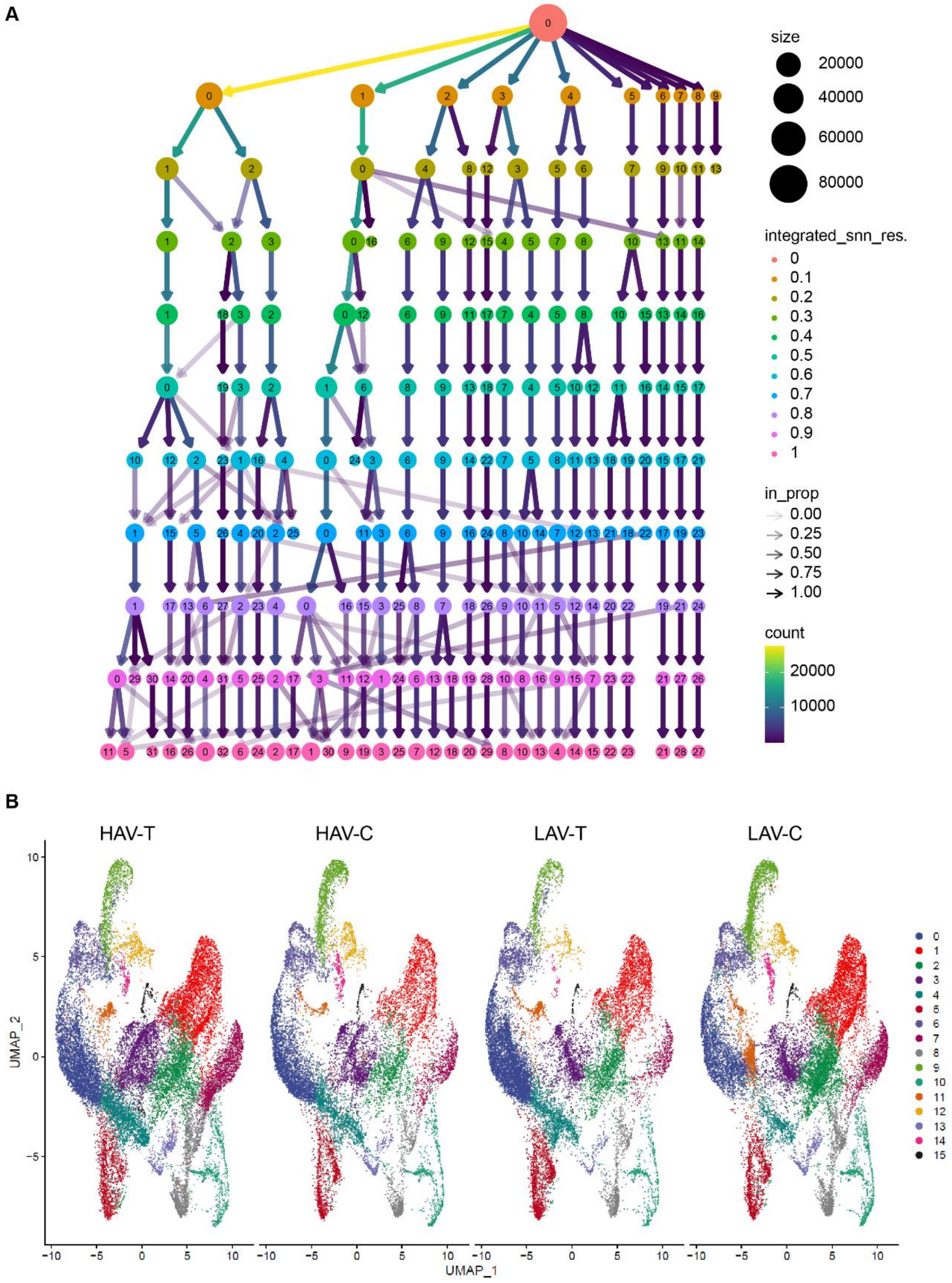


**Supplementary Figure 2. Resolution optimization and clustering analysis of integrated single-cell data from four lettuce root tip samples. A.** Clustree plot illustrating the dynamic changes of cell clusters under different resolution thresholds. **B.** UMAP plot of cell clusters after integration of four samples at a resolution of 0.3.

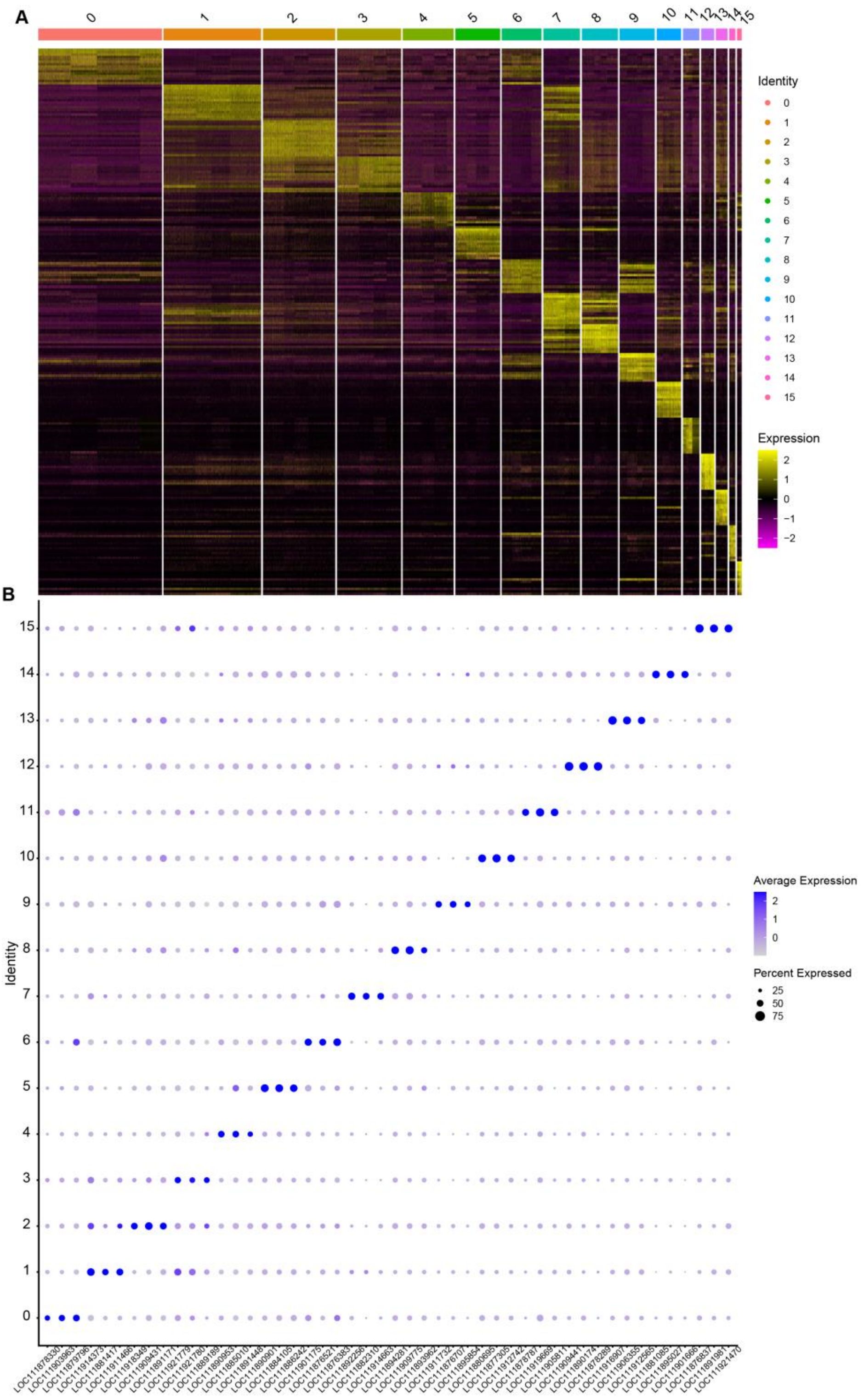


**Supplementary Figure 3. Identification and expression of cluster marker genes. A.** Heatmap showing the top 15 marker genes for each cell cluster based on the integrated single-cell data. **B.** DotPlot displaying the top 3 marker genes for each cell cluster.

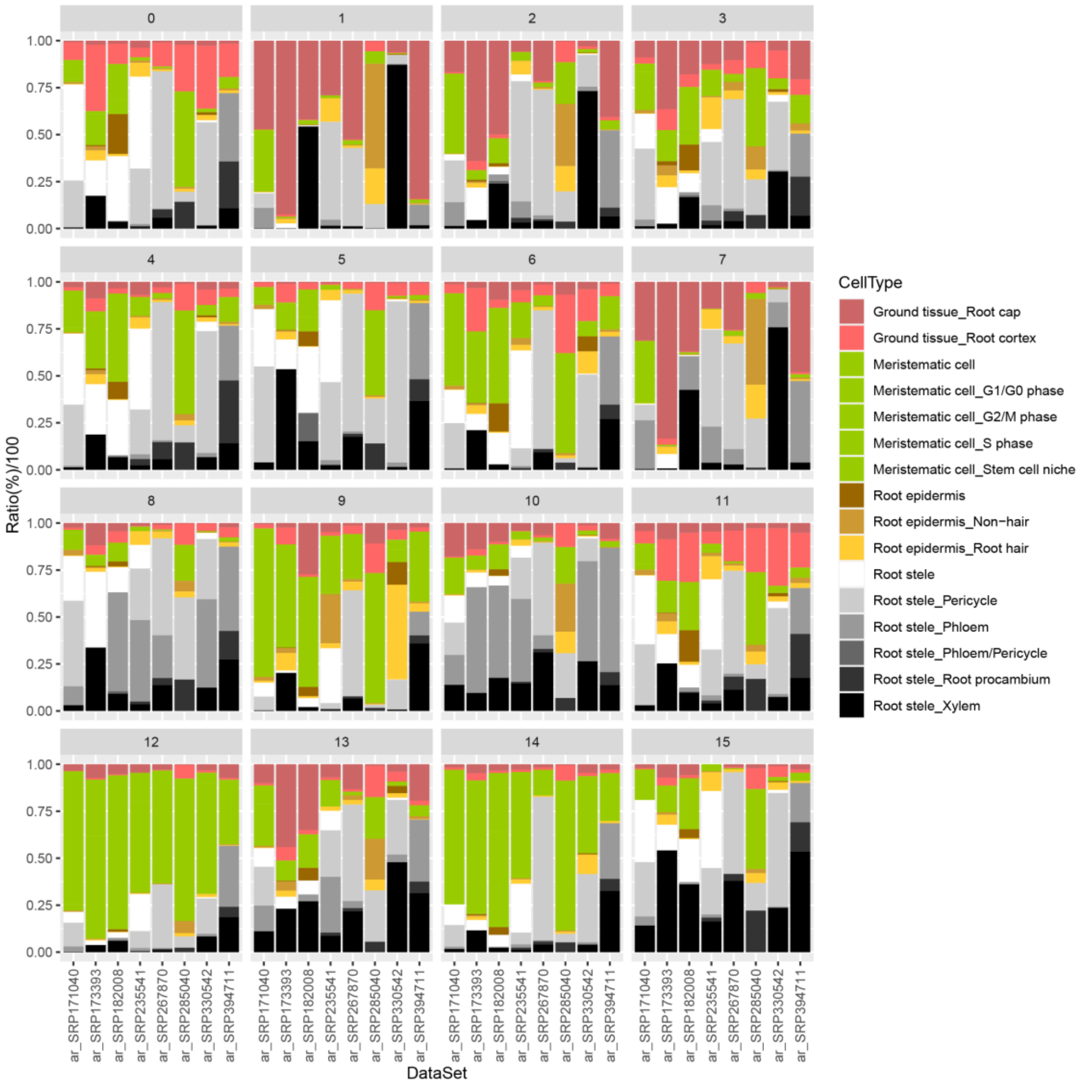


**Supplementary Figure 4. Bar plot showing the results of lettuce root tip single-cell annotation using eight Arabidopsis root tip single-cell reference datasets and the SingleR software.**

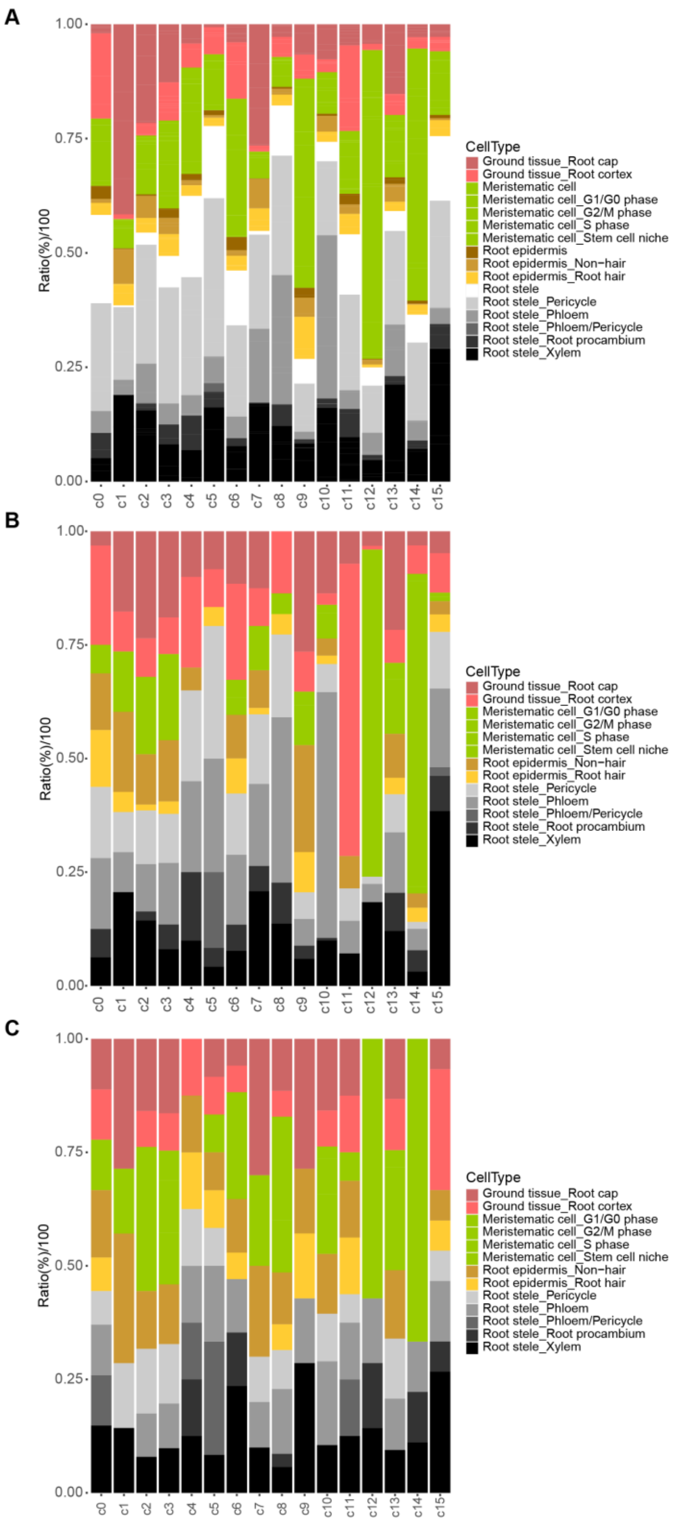


**Supplementary Figure 5. Bar plots summarizing the annotation results of lettuce root tip single-cell data based on three strategies. A.** Annotation using Arabidopsis root tip single-cell reference datasets (summarized from Supplementary Figure 4). **B.** Annotation based on alignment with Arabidopsis marker genes. **C.** Annotation guided by GO enrichment analysis matching.

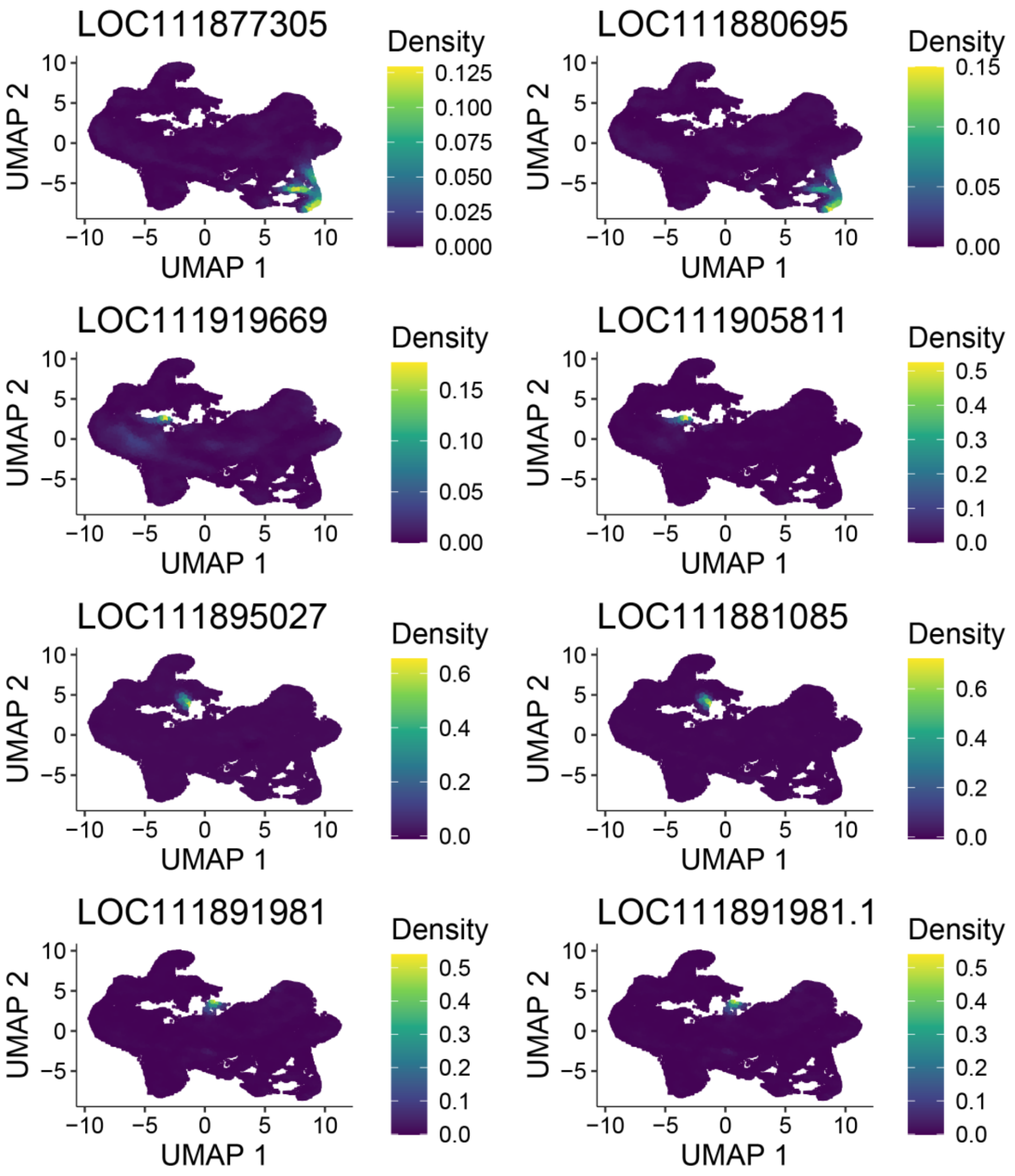


**Supplementary Figure 6. FeaturePlot visualization of cell type-specific marker genes.** The figure shows the expression distribution of representative marker genes for each cell cluster projected onto the UMAP dimensionality reduction space. Each gene is associated with its specific cell type and cluster ID:

Phloem (c10): LOC111877305, LOC111880695;

Procambium (c11): LOC111919669, LOC111905811;

Meristematic cells (c14): LOC111895027, LOC111881085;

Xylem (c15): LOC111891981, LOC111921470.

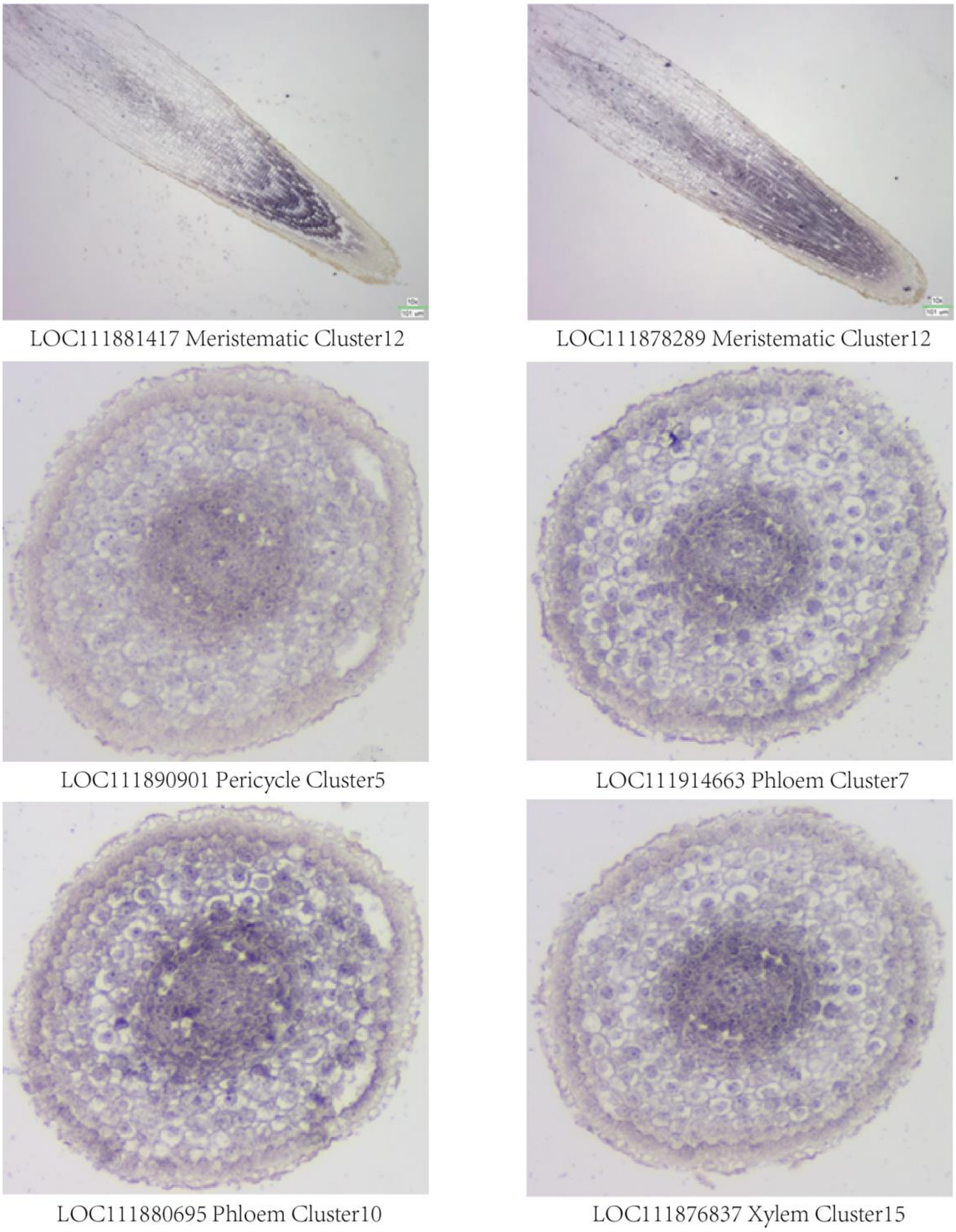


**Supplementary Figure 7. RNA fluorescence in situ hybridization (RNA FISH) validation of selected cell type-specific marker genes in lettuce root tips.**

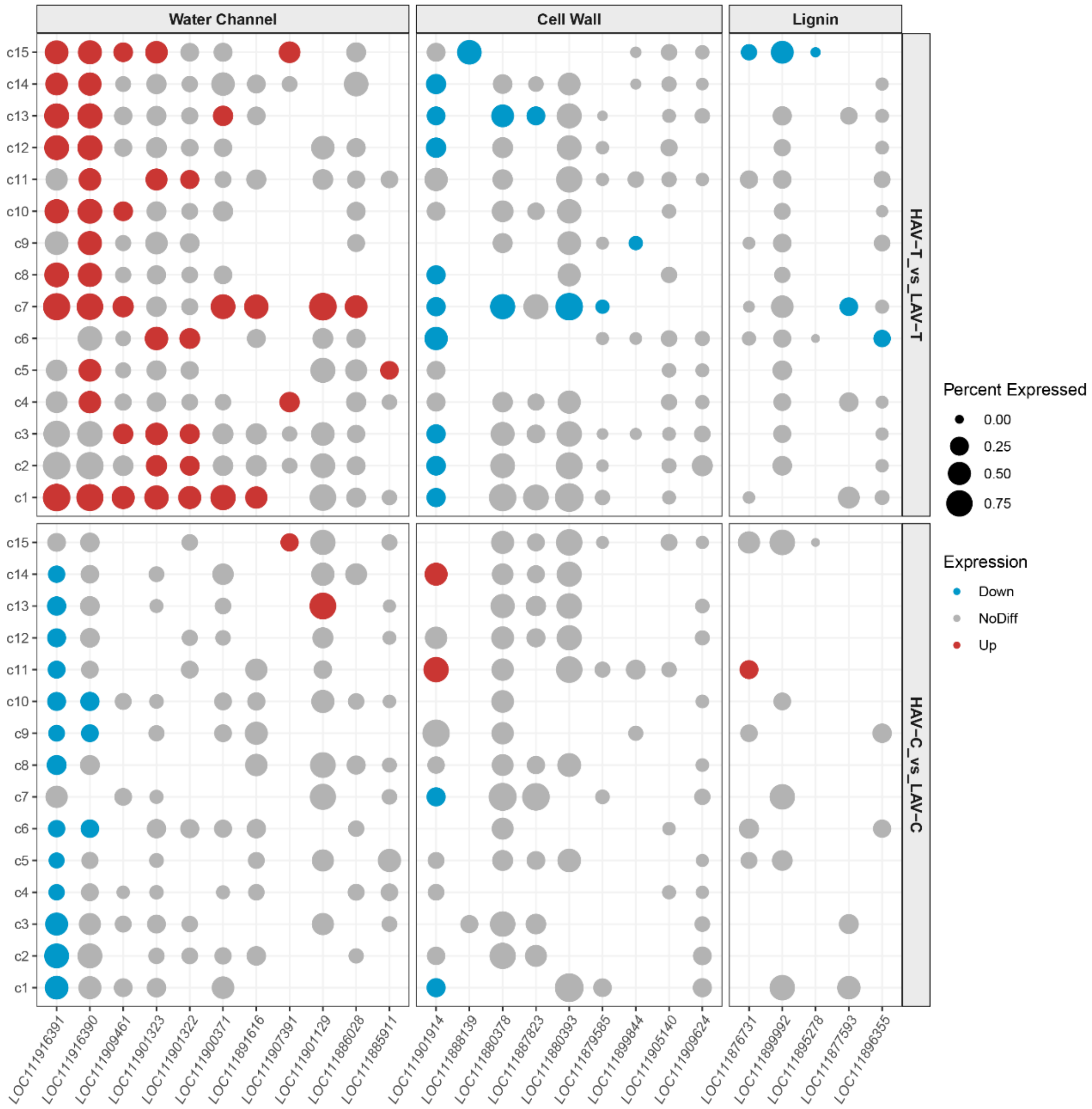


**Supplementary Figure 8. DotPlot visualization of water-channel, cell-wall, and lignin biosynthesis gene expression across cell clusters, with comparisons between treatment groups.**

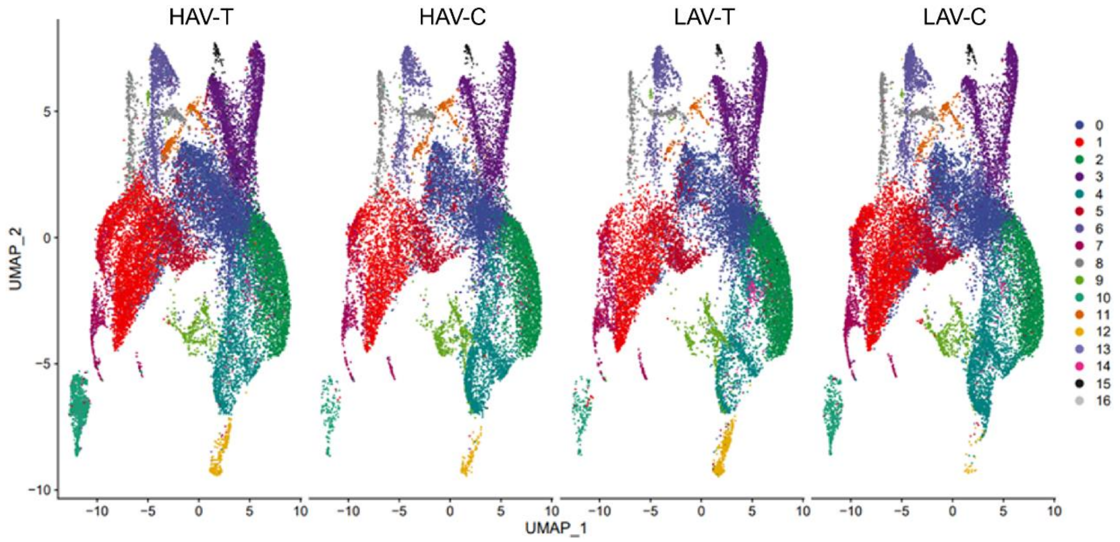


**Supplementary Figure 9. UMAP plot of HIT-scISOseq cell clusters after integration of four samples.**

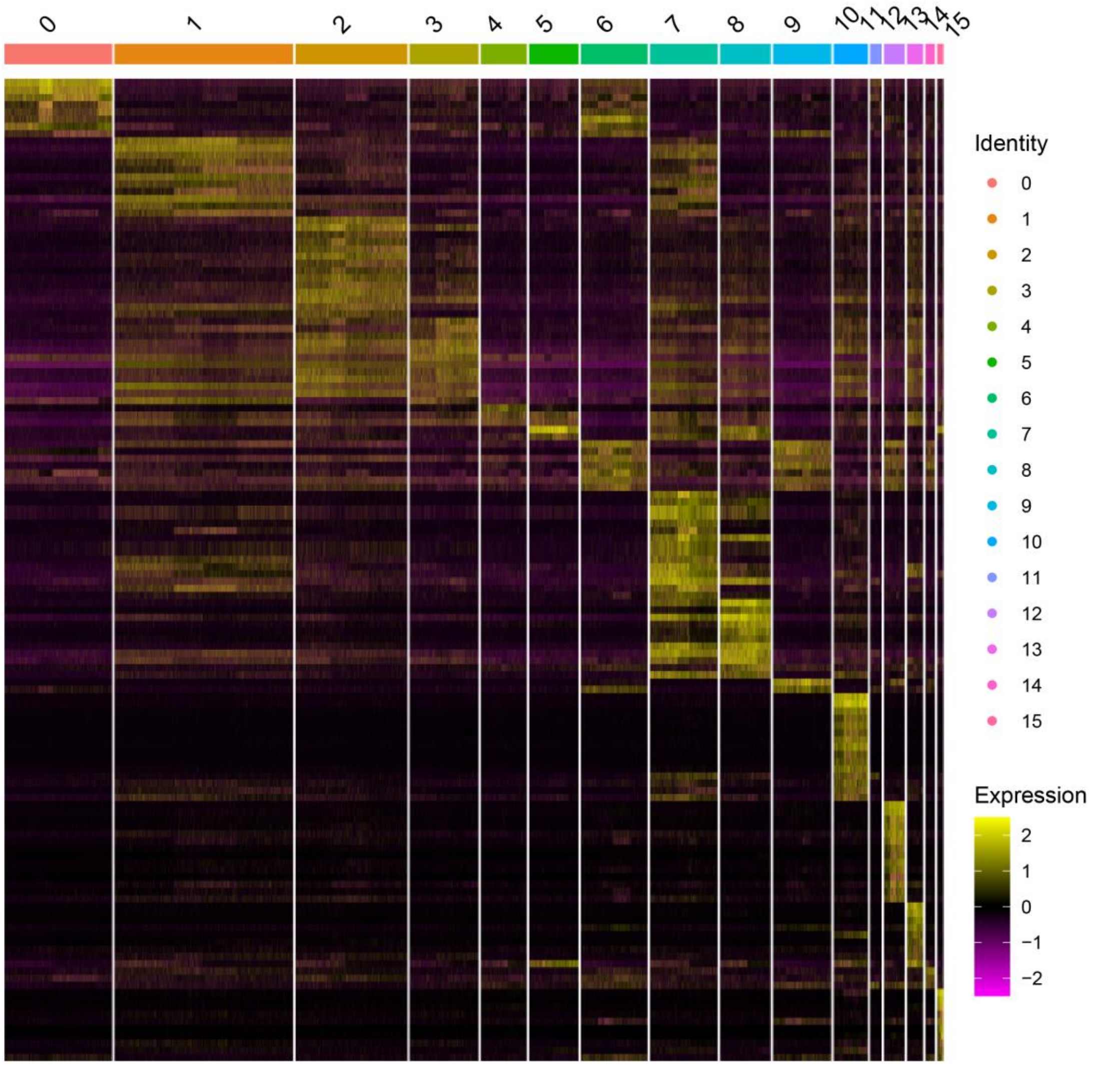


**Supplementary Figure 10. Heatmap showing the top 15 marker isoform for each cell cluster based on the integrated single-cell data.**

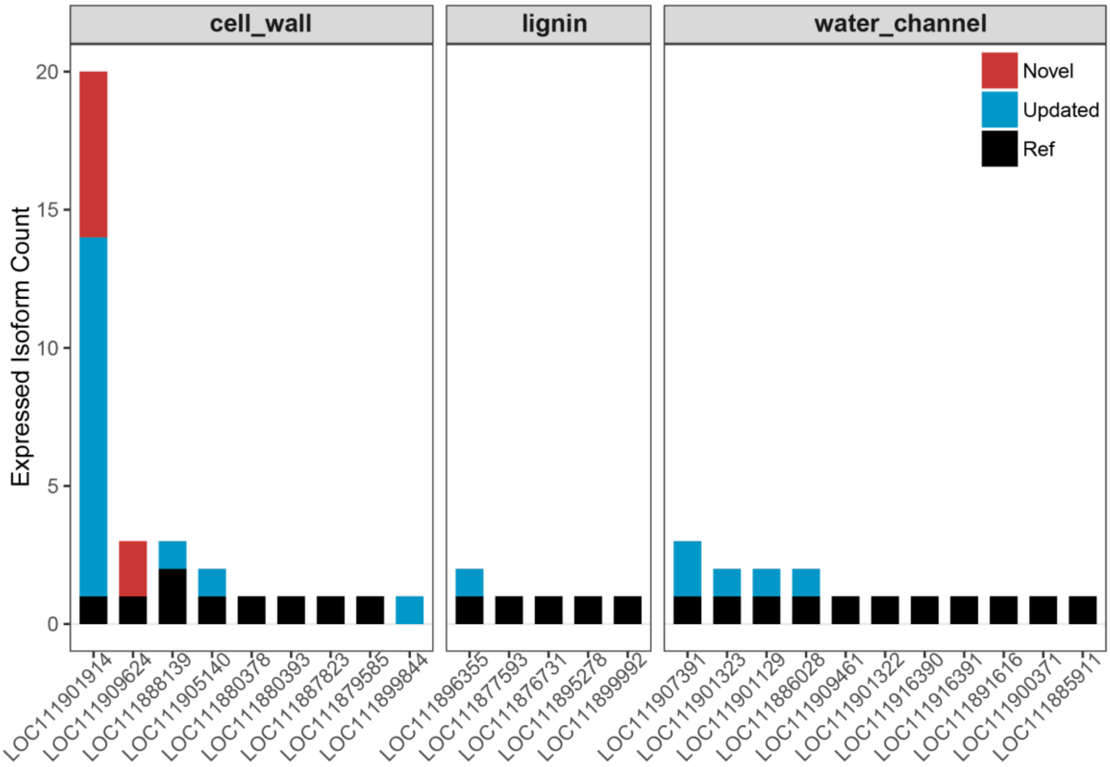


**Supplementary Figure 11. Expression profiles of RNA isoforms from genes associated with cell wall synthesis, lignin synthesis, and aquaporin activity in the HIT-scISOseq dataset. Transcripts are categorized as "Ref" (from NCBI annotation), "Updated" (added in our previous annotation upgrade), and "Novel" (newly identified in this study).**

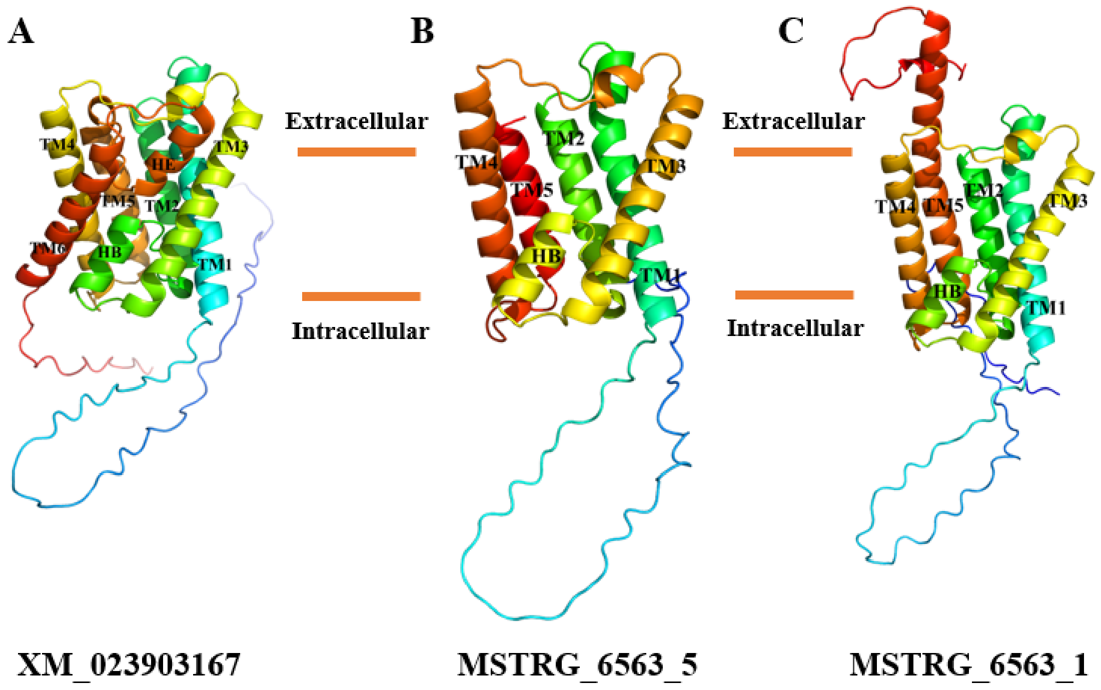


**Supplementary Figure 12. Predicted three-dimensional structures of three proteins encoded by the aquaporin gene LOC111907391.** A-C. Overall structures viewed parallel to the membrane plane: (**A**) protein encoded by the full-length transcript rna-XM_023903167; (**B**) protein encoded by the truncated transcript MSTRG_6563_5; (**C**) protein encoded by the truncated transcript MSTRG_6563_1.

#### ■ Supplementary Tables

**Supplementary Table 1.** Statistics of PFOA accumulation in two lettuce varieties.

| | PFOA concentration (0.1 mg/L) | | PFOA concentration (0.5 mg/L) | |
|---|---|---|---|---|
| The content of PFOA in the aboveground part (ug/g) | HAV | LAV | HAV | LAV |
| Rep1 | 0.264828 | 0.140641 | 2.808481 | 1.193855 |
| Rep2 | 0.321350 | 0.135629 | 2.649481 | 1.427734 |
| Rep3 | 0.737497 | 0.121771 | 2.731193 | 1.173057 |
| Mean | 0.441225 | 0.132680 | 2.729719 | 1.264882 |

**Supplementary Table 2.** Statistics of single-cell NGS sequencing data.

| | HAV-T | | HAV-C | | LAV-T | | LAV-C | |
|---|---|---|---|---|---|---|---|---|
| | update | noupdate | update | noupdate | update | noupdate | update | noupdate |
| CellRanger | | | | | | | | |
| Estimated Number of Cells | 29804 | 31,723 | 21599 | 21342 | 23746 | 25,493 | 26916 | 26826 |
| Fraction Reads in Cells | 73.20% | 80.70% | 61.40% | 65.10% | 60.80% | 69.00% | 64.30% | 69.00% |
| Mean Reads per Cell | 1495 | 16437 | 23542 | 23825 | 21151 | 19702 | 25458 | 25543 |
| Median Genes per Cell | 1182 | 1036 | 791 | 763 | 870 | 19702 | 873 | 826 |
| Total Genes Detected | 31429 | 29974 | 30727 | 29383 | 31196 | 29788 | 31288 | 29790 |
| Median UMI Counts per Cell | 2854 | 1896 | 2433 | 1598 | 2310 | 1447 | 2274 | 1719 |
| Raw Sequencing Data | | | | | | | | |
| Number of Reads (million) | 52.14 | 52.14 | 50.85 | 50.85 | 50.23 | 50.23 | 68.52 | 68.52 |
| Number of Skipped Reads | 0 | 0 | 0 | 0 | 0 | 0 | 0 | 0 |
| Valid Barcodes | 94.00% | 94.00% | 89.50% | 89.50% | 89.20% | 89.20% | 89.90% | 89.90% |
| Valid UMIs | 100.00% | 100.00% | 100.00% | 100.00% | 99.90% | 99.90% | 99.80% | 99.80% |
| Sequencing Saturation | 32.70% | 32.50% | 19.70% | 18.70% | 22.50% | 21.60% | 45.40% | 45.10% |
| Q30 Bases in Barcode | 95.70% | 95.70% | 95.70% | 95.70% | 95.90% | 95.90% | 95.30% | 95.30% |
| Q30 Bases in RNA Read | 89.10% | 89.10% | 88.00% | 88.00% | 88.50% | 88.50% | 86.50% | 86.50% |
| Q30 Bases in UMI | 95.20% | 95.20% | 95.20% | 95.20% | 95.40% | 95.40% | 94.50% | 94.50% |
| Genome Mapping | | | | | | | | |
| Mapped | 79.10% | 79.20% | 69.30% | 69.30% | 66.60% | 66.60% | 68.70% | 68.80% |
| Mapped to Genome | 73.50% | 64.90% | 63.80% | 58.90% | 61.40% | 53.60% | 63.40% | 58.50% |
| Mapped to Intergenic Regions | 4.40% | 19.30% | 4.50% | 24.20% | 4.00% | 19.80% | 4.80% | 23.00% |
| Mapped to Intronic Regions | 5.50% | 1.00% | 5.80% | 0.50% | 4.70% | 0.60% | 5.70% | 0.80% |
| Mapped to Exonic Regions | 63.60% | 44.70% | 53.50% | 34.20% | 52.70% | 33.20% | 52.90% | 34.80% |
| Mapped to Transcriptome | 52.90% | 39.90% | 38.70% | 29.10% | 40.20% | 28.40% | 39.50% | 30.10% |
| Mapped Antisense to Gene | 0.40% | 0.30% | 0.50% | 0.30% | 0.60% | 0.40% | 0.40% | 0.30% |

**Supplementary Table 3.** Statistics on the quality control of expression matrices from NGS sequencing.

| | HAV-T | HAV-C | LAV-T | LAV-C |
|---|---|---|---|---|
| Quality control conditions | | | | |
| UMI | 1200-14000 | 1100-15000 | 1200-13000 | 1200-10000 |
| Feature | 450-4500 | 320-4500 | 400-4000 | 350-3500 |
| Mitochondrion | 0-35 | 0-35 | 0-35 | 0-30 |
| Chloroplast | 0-1 | 0-1 | 0-1 | 0-1 |
| Quality control result | | | | |
| Raw Cells | 29804 | 21599 | 23746 | 26916 |
| Final Cells | 24323 | 16566 | 18372 | 21588 |

**Supplementary Table 4.** Detection statistics of RNA molecules and isoforms in NGS matrix.

| | HAV-T | HAV-C | LAV-T | LAV-C |
|---|---|---|---|---|
| Number of cell | 24323 | 16566 | 18372 | 21588 |
| Mean UMI per cell | 4098.12 | 3808.28 | 3390.33 | 3145.10 |
| Median UMI per cell | 2969 | 2258.5 | 2258 | 2447 |
| Mean genes per cell | 1712.02 | 1440.18 | 1367.89 | 1306.47 |
| Median genes per cell | 1334 | 909 | 983 | 1035 |
| Total genes detected | 27485 | 26977 | 27290 | 27267 |

**Supplementary Table 5.** Statistics of HIT-scISOseq sequencing data from lettuce root tips.

| | HAV-T | HAV-C | LAV-T | LAV-C |
|---|---|---|---|---|
| CCS Data | | | | |
| CCS Reads | 6057920 | 6265454 | 4957700 | 6123479 |
| CCS Yield (GB) | 29.14 | 28 | 22.28 | 25.8 |
| CCS MaxLen | 27177 | 27608 | 28133 | 27980 |
| CCS MeanLen | 4809.51 | 4469.18 | 4493.27 | 4212.54 |
| CCS N50Len | 5632 | 5094 | 5117 | 4730 |
| CCS Mean Passes | 16 | 15 | 23 | 20 |
| CCS MeanQV | 0.95 | 0.95 | 0.97 | 0.96 |
| FLNC Data | | | | |
| All Paire | 26816438 | 24945480 | 21778955 | 26257915 |
| FL | 23966047 | 21744782 | 20123530 | 23563227 |
| NFL | 741822 | 760862 | 379045 | 615674 |
| Unknow | 2108569 | 2439836 | 1276380 | 2079014 |
| FL(%) | 89.37 | 87.17 | 92.40 | 89.74 |
| NFL(%) | 2.77 | 3.05 | 1.74 | 2.34 |
| Unknow(%) | 7.86 | 9.78 | 5.86 | 7.92 |
| FL MeanLen | 628.96 | 590.97 | 675.60 | 560.64 |
| FL N50 | 789 | 809 | 835 | 738 |

**Supplementary Table 6.** Detection statistics of RNA molecules and isoforms in HIT-scISOseq matrix.

| | HAV-T | HAV-C | LAV-T | LAV-C |
|---|---|---|---|---|
| Number of cell | 24323 | 16566 | 18372 | 21588 |
| Mean UMI per cell | 350.13 | 314.16 | 306.44 | 290.64 |
| Median UMI per cell | 229 | 161 | 179 | 192 |
| Mean isoforms per cell | 281.93 | 244.14 | 241.99 | 229.35 |
| Median isoforms per cell | 197 | 135 | 155 | 162 |
| Total isoforms detected | 35888 | 32120 | 32073 | 34039 |

**Supplementary Table 7.** Statistics of cell numbers in each cell cluster.

| | HAV-T | HAV-C | LAV-T | LAV-C |
|---|---|---|---|---|
| Cluster0 | 3779 | 3192 | 5107 | 2615 |
| Cluster1 | 3838 | 1893 | 2253 | 3661 |
| Cluster2 | 2528 | 1248 | 1683 | 3192 |
| Cluster3 | 2650 | 1742 | 1500 | 1754 |
| Cluster4 | 2281 | 1437 | 1547 | 808 |
| Cluster5 | 1362 | 1114 | 1545 | 1301 |
| Cluster6 | 1230 | 1106 | 1215 | 1219 |
| Cluster7 | 1798 | 758 | 511 | 1327 |
| Cluster8 | 1486 | 739 | 803 | 1272 |
| Cluster9 | 919 | 1196 | 611 | 1483 |
| Cluster10 | 773 | 742 | 556 | 865 |
| Cluster11 | 398 | 228 | 480 | 880 |
| Cluster12 | 465 | 458 | 205 | 432 |
| Cluster13 | 453 | 347 | 204 | 380 |
| Cluster14 | 173 | 241 | 60 | 258 |
| Cluster15 | 190 | 125 | 92 | 141 |

**Supplementary Table 8.** scPlantDB datasets used for cell-type annotation in lettuce root tip.

| DataSet | Species | Tissue | Author | Published Date | Journal |
|---|---|---|---|---|---|
| SRP171040 | Arabidopsis thaliana | Root tip | Ryu KH et al. | 2019 | Plant Physiol |
| SRP173393 | Arabidopsis thaliana | Root tip | Denyer T et al. | 2019 | Dev Cell |
| SRP182008 | Arabidopsis thaliana | Root tip | Tian-Qi Zhang et al. | 2019 | Mol Plant |
| SRP235541 | Arabidopsis thaliana | Root tip | Wendrich JR et al. | 2020 | Science |
| SRP267870 | Arabidopsis thaliana | Root tip | Shahan R et al. | 2022 | Dev Cell |
| SRP285040 | Arabidopsis thaliana | Root tip | Yanping Long et al. | 2021 | Genome Biol |
| SRP330542 | Arabidopsis thaliana | Root tip | Moritz Graeff et al. | 2021 | Mol Plant |
| SRP394711 | Arabidopsis thaliana | Root tip | Nolan TM | 2023 | Science |

**Supplementary Table 9.** Homologs of conserved Arabidopsis marker genes in lettuce and their associated cell types.

| Arabidopsis GeneID | Arabidopsis Gene Name | Arabidopsis Cell-Type | Lettuce GeneID | Lettuce Cell-Type |
|---|---|---|---|---|
| AT5G57620 | AtMYB36 | endodermal | LOC111879582 | Cluster1,Cluster11 |
| AT5G49270 | COBL9 | root hair | LOC111915373 | Cluster1,Cluster2 |
| AT5G57620 | AtMYB36 | endodermal | LOC111909570 | Cluster2,Cluster6 |
| AT1G22710 | SUC2 | stele | LOC111911847 | Cluster7,Cluster8,Cluster13 |
| AT3G61470 | LHCA2 | lateral root | LOC111919349 | Cluster7 |
| AT4G19840 | ATPP2-A1 | phloem | LOC111902900 | Cluster8 |
| AT3G20840 | PLT1 | meristematic | LOC111877456 | Cluster9,Cluster12 |
| AT3G54890 | LHCA1 | lateral root | LOC111877237 | Cluster9,Cluster12 |
| AT1G79580 | ANAC033 | root cap | LOC111890519 | Cluster9 |
| AT4G19840 | ATPP2-A1 | phloem | LOC111902904 | Cluster10 |
| AT5G49270 | COBL9 | root hair | LOC111914140 | Cluster14 |
| AT2G37090 | IRX9 | xylem | LOC111915198 | Cluster15 |

**Supplementary Table 10.** RNA FISH probe sequence.

| Cluster | GeneID | Probe sequence | GC | Tm |
|---|---|---|---|---|
| Cluster1 | LOC111881417 | CGTTAGCCGTTCGATCAACGCTTTGGTGCA | 53% | 67℃ |
| Cluster5 | LOC111890901 | CCAAACAACAACGCTGTTGATGCTGAAGGT | 47% | 65℃ |
| Cluster7 | LOC111914663 | TTGACCAGCTGTTCCACCTGCATATACAGG | 50% | 65℃ |
| Cluster10 | LOC111880695 | CAGAAGTTCAACACCATAATGAGCGACGCC | 50% | 64℃ |
| Cluster12 | LOC111878289 | GCCTCTGAAATGGAAGCTTACGGATCAGGA | 50% | 63℃ |
| Cluster15 | LOC111876837 | TTCGTTGGACTCGGTTTGTTTCAACTCAGG | 47% | 64℃ |

**Supplementary Table 11.** Differentially expressed genes in distinct states along the developmental trajectory.

| Stage | DEG Number | Examples of DEGs |
|---|---|---|
| State1 | 1274 | LOC111913912, LOC111896763, LOC111902212 |
| State2 | 1743 | LOC111892700, LOC111906274, LOC111918372 |
| State3 | 1685 | LOC111909441, LOC111887456, LOC111905577 |
| State4 | 989 | LOC111901140, LOC111901128, LOC111880347 |
| State5 | 1845 | LOC111908934, LOC111918248, LOC111906382 |

**Supplementary Table 12.** Differential gene expression results among lettuce root tip cell clusters.

| | HAV-T Up | LAV-T Up |
|---|---|---|
| Cluster1 | 227 | 250 |
| Cluster2 | 115 | 86 |
| Cluster3 | 107 | 100 |
| Cluster4 | 82 | 61 |
| Cluster5 | 90 | 42 |
| Cluster6 | 87 | 149 |
| Cluster7 | 276 | 275 |
| Cluster8 | 112 | 42 |
| Cluster9 | 98 | 161 |
| Cluster10 | 162 | 105 |
| Cluster11 | 127 | 73 |
| Cluster12 | 96 | 90 |
| Cluster13 | 210 | 149 |
| Cluster14 | 105 | 86 |
| Cluster15 | 169 | 115 |

**Supplementary Table 13.** Transcript expression statistics of key genes involved in PFOA accumulation.

| GeneID | GO term | Gene description | Annotation | Expressed |
|---|---|---|---|---|
| LOC111909624 | cell wall organization | probable xyloglucan endotransglucosylase/hydrolase protein 23 | 3 | 3 |
| LOC111880378 | cell wall organization | CASP-like protein 1 | 1 | 1 |
| LOC111880393 | cell wall organization | CASP-like protein 1 | 1 | 1 |
| LOC111887823 | cell wall macromolecule catabolic process | endochitinase EP3 | 1 | 1 |
| LOC111879585 | cell wall organization | probable xyloglucan endotransglucosylase/hydrolase protein 23 | 1 | 1 |
| LOC111901914 | structural constituent of cell wall | extensin-1 | 24 | 20 |
| LOC111899844 | structural constituent of cell wall | extensin-2-like | 3 | 1 |
| LOC111888139 | cell wall macromolecule catabolic process | chitinase-like protein 2 | 6 | 3 |
| LOC111905140 | cell wall organization | xyloglucan endotransglucosylase protein 1 | 3 | 2 |
| LOC111877593 | lignin biosynthetic process | caffeoyl-CoA O-methyltransferase 5 | 1 | 1 |
| LOC111896355 | lignin biosynthetic process | probable cinnamyl alcohol dehydrogenase | 3 | 2 |
| LOC111876731 | lignin biosynthetic process | caffeoyl-CoA O-methyltransferase | 1 | 1 |
| LOC111895278 | lignin biosynthetic process | caffeoyl-CoA O-methyltransferase | 1 | 1 |
| LOC111899992 | lignin biosynthetic process | probable cinnamyl alcohol dehydrogenase 1 | 3 | 1 |
| LOC111909461 | aquaporin TIP | aquaporin TIP1-2 | 1 | 1 |
| LOC111901322 | aquaporin PIP | aquaporin PIP1-1 | 1 | 1 |
| LOC111901323 | aquaporin PIP | aquaporin PIP1-1 | 7 | 2 |
| LOC111916390 | aquaporin PIP | aquaporin PIP2-4 | 1 | 1 |
| LOC111916391 | aquaporin PIP | aquaporin PIP2-4 | 1 | 1 |
| LOC111901129 | aquaporin PIP | aquaporin PIP1-3 | 5 | 2 |
| LOC111891616 | aquaporin TIP | probable aquaporin TIP-type RB7-5A | 1 | 1 |
| LOC111900371 | aquaporin PIP | aquaporin PIP1-3 | 1 | 1 |
| LOC111886028 | aquaporin PIP | aquaporin PIP2-2 | 7 | 2 |
| LOC111885911 | aquaporin TIP | probable aquaporin TIP-type | 1 | 1 |
| LOC111907391 | water channel activity | probable aquaporin NIP5-1 | 6 | 3 |
